# Recurrent chromosome destabilization through repeat-mediated rearrangements in a fungal pathogen

**DOI:** 10.1101/2023.07.14.549097

**Authors:** Simone Fouché, Ursula Oggenfuss, Bruce A. McDonald, Daniel Croll

## Abstract

Genomic instability caused by chromosomal rearrangements has severe consequences for organismal fitness and progression of cancerous cell lines. The triggers of destabilized chromosomes remain poorly understood but likely co-locate with fragile sites. Here, we retrace a runaway chromosomal degeneration process observed in a fungal pathogen using telomere-to-telomere assemblies across an experimental progeny. We show that the same fragile sites triggered reproducible, large-scale rearrangements through non-allelic recombination. Across our four-generation progeny, chromosomal rearrangements were accompanied by non-disjunction events leading to aneuploid progeny with up to four chromosomal copies. We identify a specific transposable element family co-locating with fragile sites, likely triggering ongoing repeated chromosomal degeneration. The element has recently been associated with lower virulence of the pathogen and has undergone an expansion of copy numbers across the genome. Chromosome sequences are also targeted by repeat-induced point mutation, a genome defense mechanism actively leading to hypermutation on duplicated sequences. Our work identifies the exact sequence triggers that initiate chromosome instability and perpetuate degenerative cycles. Dissecting proximate causes leading to runaway chromosomal degeneration could expand our understanding of chromosomal evolution beyond fungal pathogens.

## Introduction

Meiosis is a highly conserved process in eukaryotes, whereby homologous chromosomes pair, undergo recombination, and separate into daughter cells. Aberrations during the faithful transmission of chromosomes through meiosis can have serious consequences for an organism. Non-disjunction events resulting in additional or fewer chromosomal copies occur frequently in humans, and are the leading causes of miscarriage (*1*). Similarly, non-disjunction events and chromosome rearrangements occur in somatic lines, and may lead to genomic instability, which is a hallmark of cancers (*2*). However, factors that cause fragility of pre-cancerous genomes or that trigger meiotic errors are largely unknown. Chromosomal breakage is a major factor contributing to instability and occurs often at specific loci referred to as fragile sites (*3*). Though the locations of many mitotic fragile sites have been identified in the human genome (*4*), how they contribute mechanistically to breakage remains poorly understood. Fragile sites for non-allelic homologous or ectopic recombination during meiosis remain largely unknown, possibly because major rearrangements tend to be lethal and are quickly purged from the population by purifying selection. Investigations of non-lethal rearrangements offer a promising approach to unravel sequence determinants of fragile sites.

Accessory chromosomes in plant and fungal genomes are powerful models to study non-lethal chromosome rearrangements. These chromosomes (also called supernumerary or B) are present in a karyotype in addition to the regular chromosomes, contain no essential genes, and show broad presence/absence within species (*5*). It is unclear what mechanisms generate these chromosome rearrangements and whether fragile sites are involved. For example, the accessory chromosomes of *Magnaporthe oryzae* contain many rearrangements and frequent interchromosome translocations between core and accessory chromosomes (*6*). Accessory chromosomes are hypothesized to originate from core chromosomes through rearrangements involving the terminal regions of core chromosomes. *Zymoseptoria tritici*, a haploid fungal pathogen of wheat, exhibits some of the most extreme degrees of structural variation observed within fungal species, including large differences in terms of chromosome length, transposable element (TE) and gene content, and recombination rate, as well as telomere and centromere structures (*7–13*). Chromosome rearrangements occur frequently during meiosis, with higher rates of rearrangements observed in the accessory chromosomes (*7*, *14*).

We analyzed a rearranged accessory chromosome of *Z. tritici*, , which was generated repeatedly during meiosis through an unknown mechanism (*7*). Here, we show how the rearranged chromosome was generated and how further rearrangements occurred through subsequent rounds of meiosis. We also find evidence that a transposable element acts as the specific sequence trigger of the chromosome rearrangement and degenerative cycles.

## Results

### Origin of a complex chromosome rearrangement

The rearrangement of the *Z. tritici* accessory chromosome 17 was first discovered in two F1 progeny A66.2 and A2.2 (fig. 1A) (*7*). The enlarged nature of chromosome 17 in these progeny was confirmed by pulsed-field gel electrophoresis (PFGE) as well as southern hybridization (fig. 1B) (*7*). To assess how frequently rearrangements of chromosome 17 occur through meiosis, we screened an additional 48 progeny from the same cross with a segment-specific PCR assay for ∼500 bp regions of coding sequences at regular intervals along chromosome 17, and found that anomalies of chromosome 17 were most likely restricted to the two initially discovered progeny A66.2 and A2.2 (fig. 1A, supplementary fig. S1). A previous screen based on restriction site associated DNA sequencing of a further 228 progeny from this cross revealed four progeny with likely rearranged chromosome 17 (*14*). Finally, we assessed the frequency of rearranged chromosome 17 variants using whole-genome sequencing of 150 field-collected isolates matching the sampling location of the parental isolates. Based on read coverage variation, the population carries likely three disomic or partially duplicated variants of chromosome 17 (*15*).

**Figure 1:**
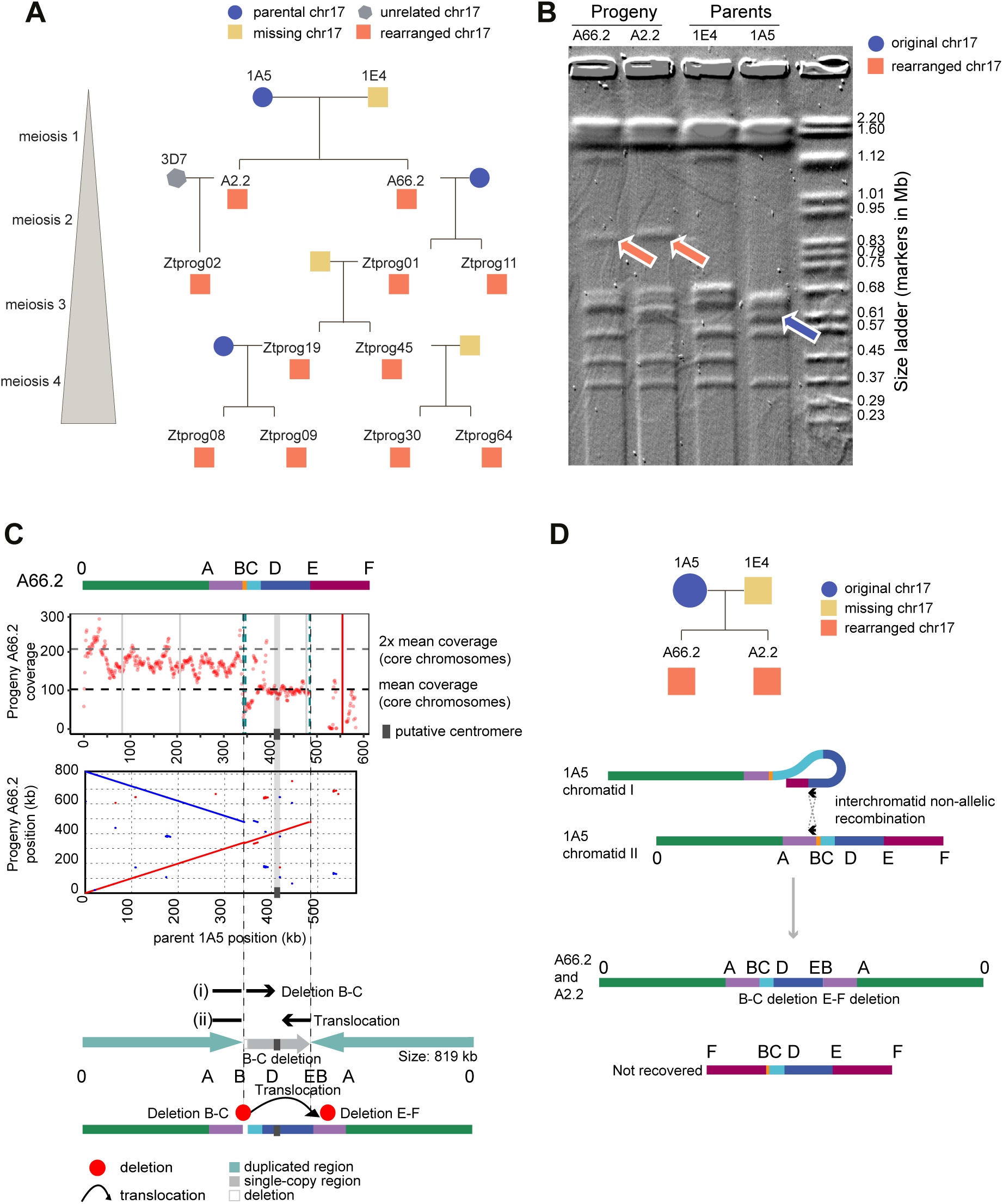
Pedigree analyses revealing repeated occurrence of enlarged chromosome 17. (A) Pedigree of four rounds of meiosis. The colors indicate whether the isolate carries a parental, unrelated (not carried by a parent), rearranged, or no chromosome 17 according to pulsed-field gel electrophoresis (PFGE) analysis. (B) PFGE of chromosomes from the progeny (A66.2 and A2.2) and the parents (1E4 and 1A5), adapted from (*7*). Orange arrows indicate the enlarged chromosome 17, the blue arrow indicates the parental chromosome 17. (C) The coverage and breakpoints of progeny A66.2 reads mapped to the parent 1A5. Black and grey horizontal dashed lines indicate mean coverage and 2x mean coverage of the core chromosomes, respectively. Red dots indicate the mean coverage in 1 kb windows (regions with excessive, >300x coverage were removed). Dashed vertical lines indicate the chromosomal breakpoints identified from mapped reads (at positions B and E, turquoise and continuing in grey). Solid vertical lines indicate the positions of loci amplified by PCR (grey: amplification; red: no amplification). Dotplot of the assembled chromosome for progeny A66.2 compared to the parental chromosome. Inverted regions are indicated in blue. (i) and (ii) show the location and continuation of split reads. At the bottom is a schematic of the resulting enlarged chromosome in the progeny. (D) Schematic representation of the breakpoints and rearrangements between the two chromatids of 1A5 that generated the enlarged chromosome 17 (0–ABCDEBA–0) recovered in progeny A66.2. The hypothetical smaller rearrangement product (FBCDEF) was not observed.

To investigate at base-pair resolution how the enlarged chromosome 17 was generated in progeny A66.2 and A2.2, we sequenced the isolates with long-read PacBio technology. The recency of the duplication creates a significant challenge for the chromosomal assembly as duplicated regions are not expected to show sequence divergence and collapse during assembly. We used a combination of coverage and split long-read mapping approaches to overcome this challenge. First, we mapped reads from the progeny to the 1A5 genome and identified regions of chromosome 17 with higher than average coverage compared to the core chromosomes (1–13). Mean read coverage transitioned sharply along chromosome 17 (fig. 1C). We found that region 0–B in progeny A66.2 and A2.2 each had approximately double the mean coverage of the core chromosomes, implying that this region was duplicated (fig. 1C, supplementary fig. S2 and S3, supplementary table S1). Further supporting evidence included a breakpoint at the end of segment 0–B, with reads showing split mapping and linking two distinct regions of the chromosome (fig. 1C, supplementary fig. S2, supplementary table S2). Reads mapping to this position were evenly split between canonically mapping across positions B–C and split reads mapping from A–B and continue in an inverted orientation from position E towards D (fig. 1C, supplementary fig. S2). Taken together, we found that the region 0–B exists twice in the progeny genome and that the duplicated sequences are connected to two distinct locations of the chromosome at C and E (fig. 1C, supplementary fig. S2 and S3).

The rearranged chromosome 17 carries a single-copy region between C and E. A lack of coverage after position E towards F suggests that this region is absent in the rearranged chromosome. We confirmed the absence of region E–F by PCR (fig. 1B, supplementary fig. S2–3). A lack of coverage between B and C indicates a deletion of this region. Using the information on coverage and reads spanning distinct chromosomal regions, the chromosome 17 of progeny A66.2 and A2.2 are composed of region 0–B, followed by region C–E, and a second copy of region 0–B in an inverted orientation (fig. 1C; supplementary fig. S2, supplementary table S2). We identified the putative centromere to be located between position 405,779 and 415,898 based on sequence homology to the canonical chromosome 17 (*8*). Therefore, the resulting enlarged chromosome 17 is expected to carry a single copy of the centromeric region (fig. 1C). Both progeny most likely inherited an identical, enlarged chromosome 17 generated during the first round of meiosis through the same non-allelic recombination between sister chromatids at locations B and E (fig 1D). The rearrangement should also have produced a second, shorter chromosomal variant. However, the shorter variant was not recovered in progeny, despite being predicted to carry a centromere. We reconstructed the sequence of the rearranged chromosome 17 to be 819 kb in length, matching the approximate length identified by PFGE (fig. 1B).

### Sustained chromosome degeneration in subsequent rounds of meiosis

We investigated the fate of the enlarged chromosome 17 in further rounds of meiosis by performing several backcrosses (fig. 2A). New variants of chromosome 17 were already generated in the second round of meiosis (fig. 2B, 3B, supplementary fig. S3). Chromosome 17 of Ztprog11 carries a duplicated region, 0– A and the second copy of 0–A is joined to position E in an inverted orientation (fig. 2B, 3B, supplementary fig. S3, S4). Ztprog01 has two copies of chromosome 17, including a small and a large variant consistent with non-disjunction. One copy lacks the segment between B and D (fig 2B, fig 3B, supplementary fig. S3, S5). During the third round of meiosis, ZtProg19 stably inherited a chromosome 17 variant (fig. 2D, fig 3C, supplementary fig. S3, S7). Based on progeny and read mapping evidence, the second progeny ZtProg45 carries multiple variants. Variant V2 may have undergone non-disjunction (2x V1 is inherited with 1x V4) or one copy of V1 is present with one copy of V2 and V3 each, indicating rearrangements at breakpoints B and D as well as non-disjunction (fig. 2C, fig. 3C, supplementary fig. S3, S6). Therefore, two small variants are present together with one large variant.

**Figure 2:**
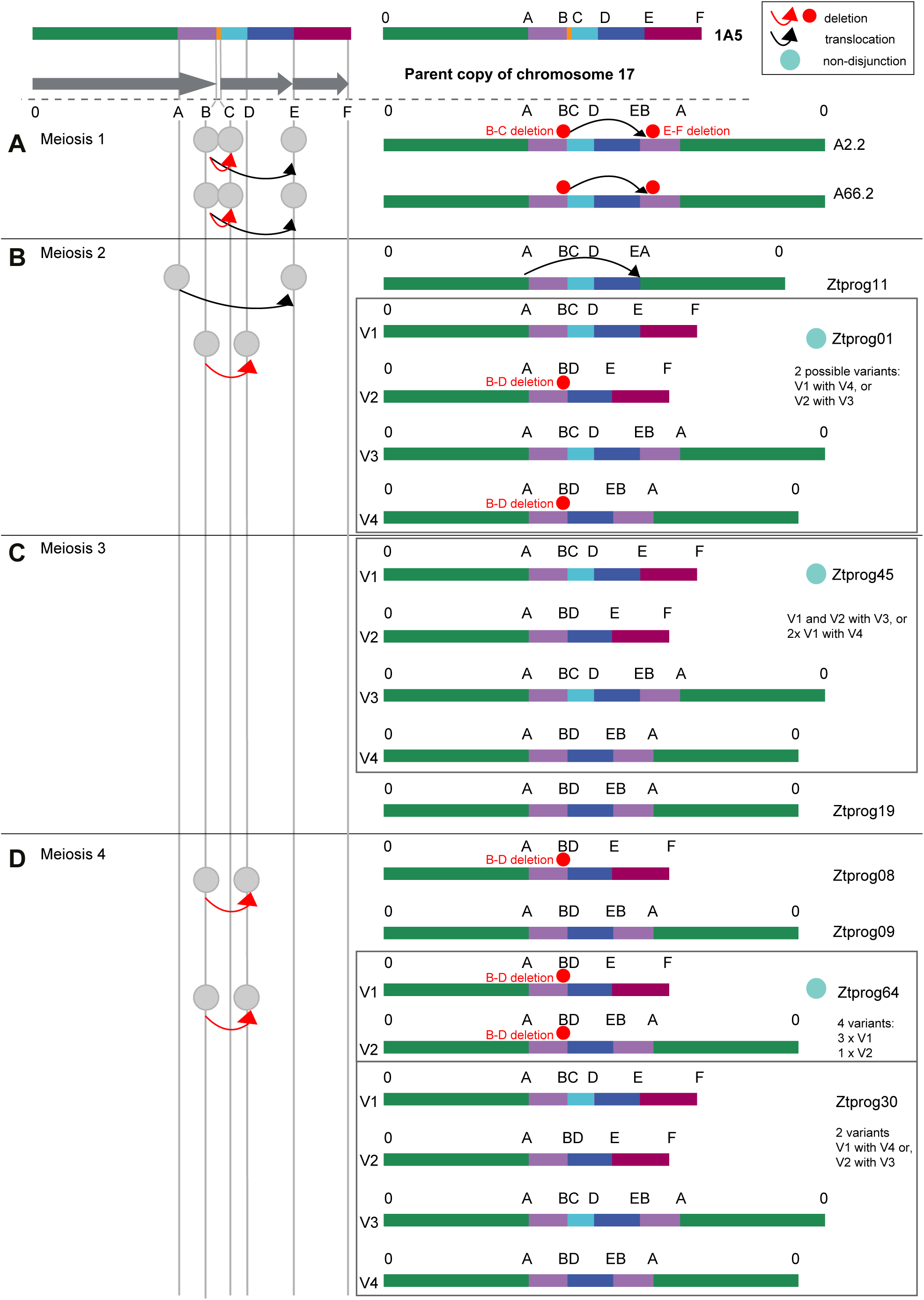
Sequence of chromosome 17 rearrangements tracked through four rounds of meiosis. Chromosome rearrangements are shown according to breakpoint positions A–F identified through split read mapping. Arrows linking different letters A–F indicate translocations and deletions for the successive rounds of meiosis. Panels on the right show schematic representations of each chromosome 17 variant present in each progeny. If multiple variants were detected, variants are labelled V1–V4.

**Figure 3:**
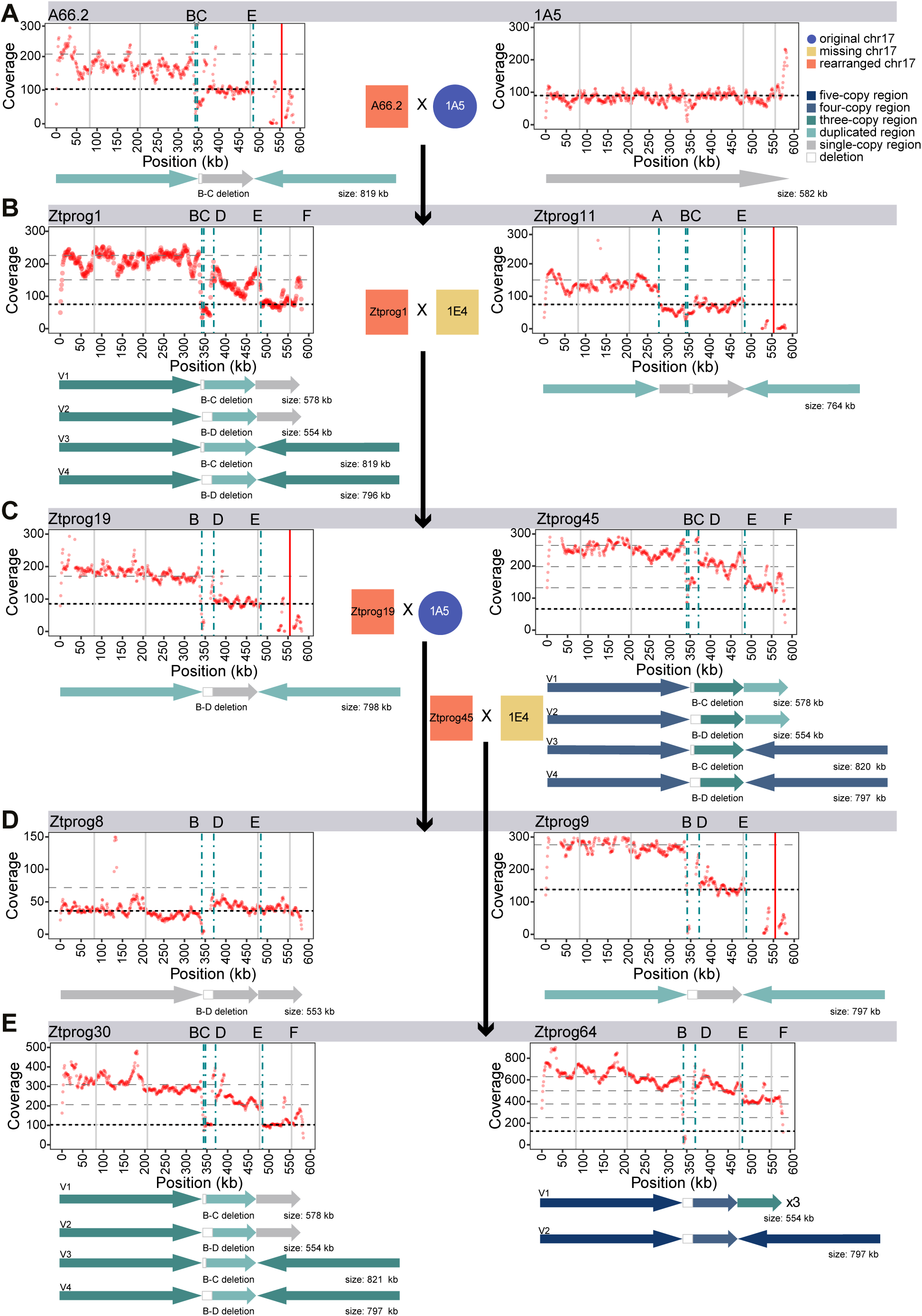
Reconstructed chromosome 17 variants based on sequence rearrangement breakpoints. For each progeny the mapped read coverage is shown relative to the 1A5 reference genome sequence. Black and grey horizontal dashed lines indicate mean coverage and 2x/3x/4x mean coverage of the core chromosomes, respectively. Red dots indicate the mean coverage in 1 kb windows. Vertical dashed lines indicate the chromosomal breakpoints A–F (see fig. 1 and 2). Solid vertical lines indicate the positions of loci amplified by PCR (grey: amplification; red: no amplification). Below each coverage plot, arrows show variants reconstructed for each progeny. If multiple variants were detected, variants are labelled with V1–V4. Variant labels are independent between progenies. Arrow color is based on the level of duplication (absent, single-copy or multiple-copy regions). The progeny is represented by (A) the progeny of the first meiotic round, (B) second, (C) third, and (D–E) fourth round.

In the last round of meiosis, the region B–D was deleted in Ztprog8 and Ztprog64 (fig. 2D, fig 3D and ii, supplementary fig. S3, S8, S9). Ztprog64 carries four chromosome 17 variants (three small and one large variant) that are missing the region B–D (fig. 2D, fig 3i and ii, supplementary fig. S3, S9). Ztprog9 most likely inherited the enlarged chromosome 17 without further rearrangements from the previous generation (fig. 2D, fig 3D, supplementary fig. S3, S10). Ztprog30 has the same chromosome complement as Ztprog01 (fig. 2D, fig 3E, supplementary fig. S3, S11). The enlarged chromosome 17 was highly unstable through further rounds of meiosis and degenerative cycles via non-allelic recombination as well as non-disjunction occurred. Rearrangements at all breakpoints except A were observed several times in the experiment.

### Degeneration of the duplicated sequence through RIP

Fungal genomes encode repeat-induced point mutations (RIP), a defense mechanism targeting duplicated sequences that dramatically reduces TE activity. We predicted that the presence of a large, duplicated region in chromosome 17 would trigger RIP (fig. 4A). To detect RIP-like mutations, we mapped reads from progeny to the parental chromosome 17 and identified regions that with transitions from G/C to A/T (fig. 4B; supplementary fig. S4). We found an overrepresentation of RIP-like mutations in region 0–B. As a control, we analyzed the rearrangement-free progeny Ztprog08 for RIP-like mutations and found indeed no overrepresentation of RIP-like mutations, indicating RIP not being active. Hence, the genomic defenses are specifically active against the new duplications appearing in degenerate chromosome 17 variants. RIP-like mutations predominantly targeted CpA dinucleotides (or TpG) (fig. 3C), matching evidence from other model fungi (*16*). Additionally, CpG dinucleotides were also frequently targeted. New RIP-like mutations were detected after a single round of meiosis, suggesting that the RIP genome defense did not yet reach saturation. RIP-like mutations never targeted the putative centromere (405,779–415,898 bp; supplementary fig. S4). Our results show that massive rearrangements and non-disjunction triggers the genome defense RIP, contributing to an increased mutation rate in the duplicated region.

**Figure 4:**
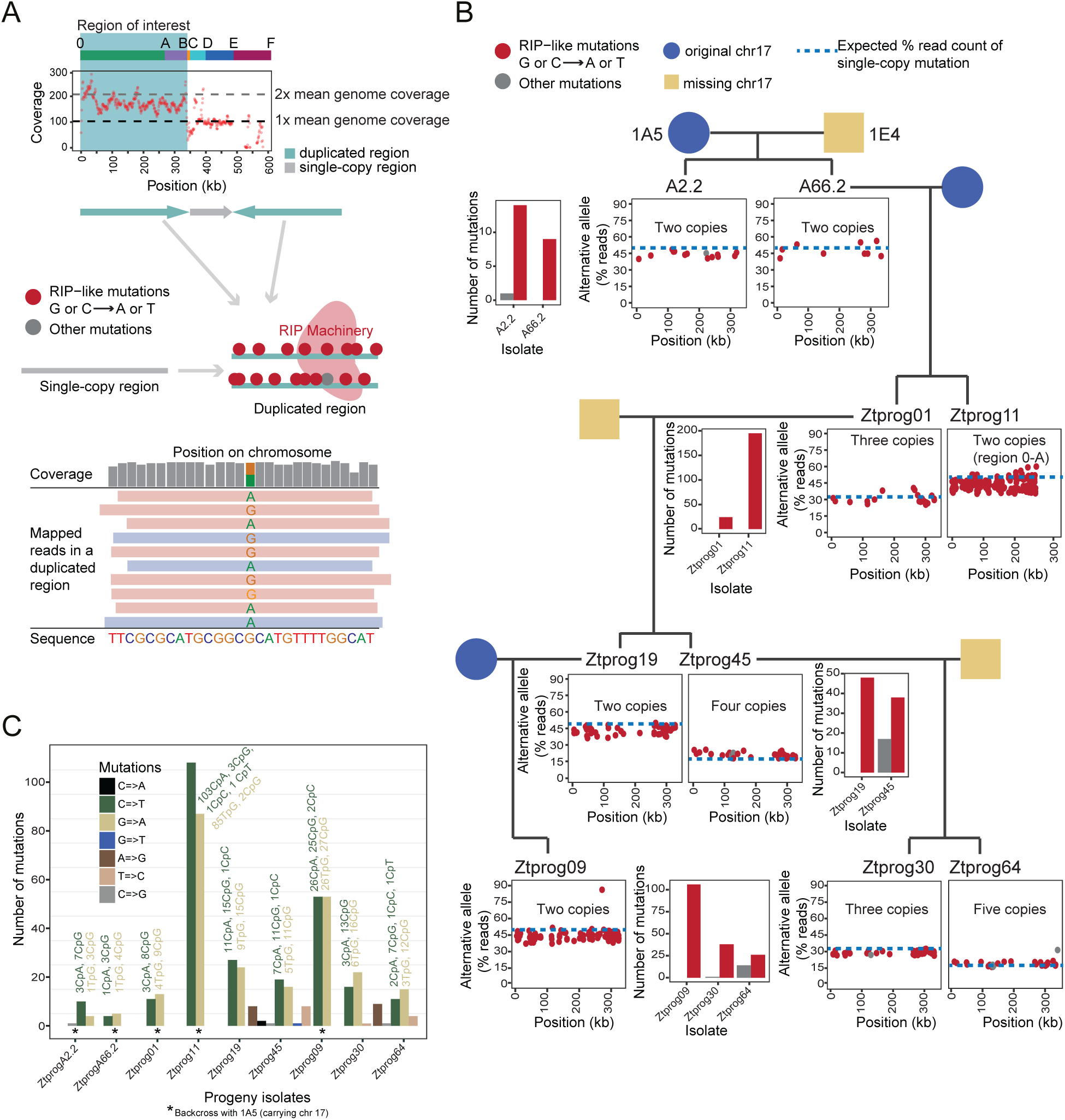
Characterization of repeat-induced point mutation (RIP) acting on duplicated sequences. (A) Read coverage along the chromosome 17 and evidence for a duplicated region 0–B (0–A in Ztprog11). The upper schematic shows how RIP mutations are introduced into duplication regions, causing an excess of Adenine and Thymine with RIP acting on Guanine or Cytosine. The lower schematic shows how mapped PacBio reads in duplicated regions were used for SNP calling, revealing potentially RIP-like mutations. (B) Overview of progeny over generations with barplots summarizing the number of RIP-like and other mutations detected per progeny generation in region 0–B (0– A in Ztprog11). Dotplots show the percentage of reads carrying the alternative allele (parent 1A5). The identified copy numbers of the region 0–B (0–A in Ztprog11) is indicated. Blue dashed lines show the expected percentage of reads, confirming a new mutation carried by at least one copy of the duplicated regions. (C) Breakdown of all mutations detected in duplicated regions of each progeny. Mutations at particular dinucleotides are summarized individually per progeny.

### Chromosomal rearrangements triggered by a recent TE burst

To understand the mechanism triggering the initial chromosome 17 rearrangement and sustaining the degeneration, we examined the breakpoint sequences. We identified a full copy of the DNA transposon *Styx* (also known as REP9) at position B and a partial copy of the same family at position C in the parental chromosome of 1A5 (fig. 5A). *Styx* was previously described as a negative regulator of virulence (*17*). Furthermore, *Styx* was shown to proliferate in the progeny and pathogen populations across continents (*18*, *19*). At position B, the element is at 582 bp from the breakpoint. A second complete copy of *Styx* was found at position D (∼8 kb in length) (fig. 5A). *Styx* has 21 copies in the 1A5 genome, including three copies on chromosome 17 (fig. 5B). Using ncbi blast, we predicted four putative coding sequences in full-length copies of *Styx*, of which one shows weak homology to RNase H and an integrase (fig. 5A) (*18*). We analyzed *Styx* copy numbers in 19 completely assembled genomes of *Z. tritici* (*13*) and in the genomes of the sister species *Z. passerinii*, *Z. brevis* and *Z. pseudotritici* (*20*) . *Styx* is present in higher copy numbers in the sister species but nearly absent in genomes sampled from the center of origin of *Z. tritici* (*18*). Our findings support a scenario of *Styx* being present in the common ancestor at low copy numbers and undergoing independent bursts in the sister species, as well as in *Z. tritici* populations in North and South America, Europe, and Australia, resulting in increased copy numbers in the parental and other isolates (fig. 5B) (*18*). The TE copy on chromosome 17 was introduced following a recent burst (fig. 5C). TE copies created during the burst are characterized by short terminal branch lengths and high GC-content (fig. 5D). The creation of the enlarged chromosome 17 was likely mediated by non-allelic homologous recombination between *Styx* copies at positions B and a different sequence with microhomology at position E. We examined the regions 1 kb up- and downstream from position A, B, and E for similar repetitive sequences. Positions B and E carry a similar 6-bp repeat that may have ultimately triggered the rearrangement (supplementary files S1– 2). Finally, near position E the repeat is 34 bp away from the breakpoint. Similarly, positions A and E had two similar 3-bp repeats (supplementary file S2 and S3).

**Figure 5:**
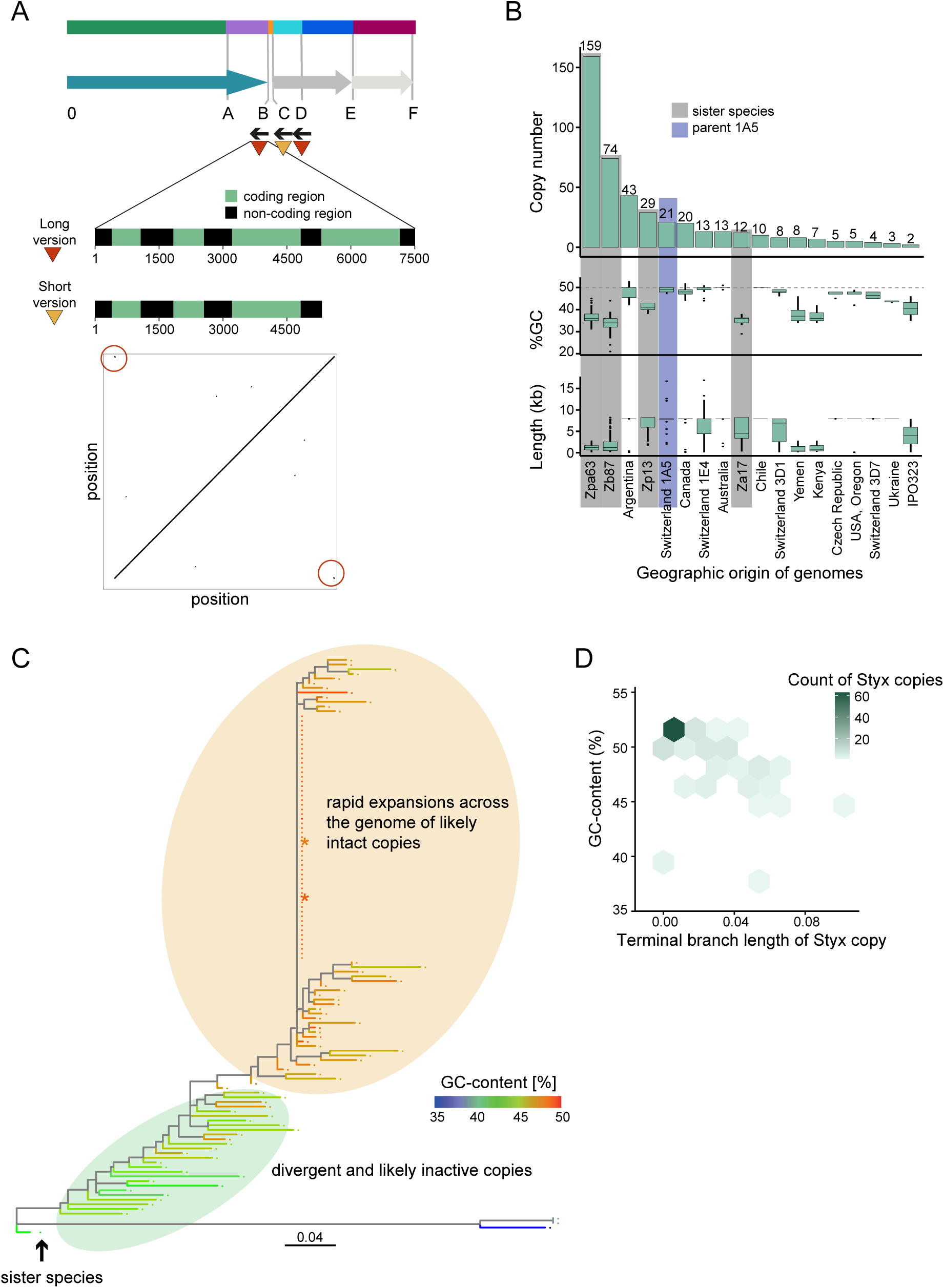
**Recent expansion of the *Styx* transposable element underpinning the chromosomal rearrangements.** (A) The length and location of coding regions of the long and short copies of *Styx*. Dotplot of the consensus sequence showing duplicated regions (red circle). (B) Copy number, GC-content, and length distribution of *Styx* copies recovered from a global panel of *Z. tritici* reference genomes. A dark grey background indicates genomes of sister species *Z. ardabiliae* (Za17), *Z. passerini* (Zpa63), *Z. brevis* (Zb87), and *Z. pseudotritici* (Zp13). (C) Phylogeny of *Styx* copies in *Z. tritici*, with *Z. pseudotritici* as an outgroup. The color scale indicates the GC content. The two full-length copies found on chromosome 17 of the parent 1A5 are indicated by a star. (D) Density plot of branch length against GC-content of individual *Styx* copies. The most recent burst of *Styx* copies show small branch lengths and high GC-content.

## Discussion

We reveal the dynamics of an unusual chromosomal rearrangement in a fungal plant pathogen. Using split long-reads, we identified the exact breakpoints and retraced the degeneration through four rounds of meiosis. We propose that the primary degenerative rearrangement was caused by non-allelic recombination between a TE and a region with microhomology to the TE. The degenerated chromosome was composed of a large, duplicated, and inverted region connected to a single-copy chromosomal segment near the centromere. The sequences serving as triggers for the rearrangement in the first round of meiosis was likely the transposable element *Styx*, being present in multiple copies on chromosome 17 and regions and sharing microhomology. As an accessory chromosome without known function, chromosome 17 is likely under relaxed selection, and chromosomal rearrangements are tolerated. However, chromosomal rearrangements co-locating with *Styx* copies were also detected in core chromosomes of the same cross (*18*). Subsequent rounds of meiosis increased the spectrum of rearrangement breakpoints consistent with runaway chromosomal degeneration. Chromosomal segments affected by the rearrangement were repeatedly affected multiple times and three of these segments co-localize with copies of *Styx*. Concurrent with the degeneration, non-disjunction events of chromosome 17 increased along the progeny, characterized by disomy and trisomy. The frequency of non-disjunction events increased with the presence of chromosome 17 variants in both parents, suggesting that opportunities for mispairing of chromatids increases the likelihood of aberrant segregation. The degenerating chromosome was furthermore affected by the genomic defense mechanism RIP, introducing mutations into recently duplicated sequences. RIP on very large duplicated sequences might have a titration effect, leading to less dense RIP mutations in smaller duplicated sequences (*21*). Overall, the identity of the paired parental genotypes had a major influence on the likelihood of rearrangements observed in the progeny.

The copy-number amplification in progeny Ztprog64 and Ztprog30 shares hallmarks of breakage-fusion-bridge (BFB) cycles observed in many cancer lines (*22–27*). Following the initial degeneration, chromosome 17 was stabilized in pairings of a single rearranged chromosome variant with a parental chromosomal variant (*i.e.,* Ztprog8 and Ztprog9). This stabilization suggests that the progression of the degenerative cycle was interrupted through proper segregation. The chromosome 17 variant in Ztprog8 was generated through non-allelic homologous recombination between two *Styx* copies. The repair of chromosomes through recombination between repeats is known to produce intermediary chromosomes that can ultimately produce stable karyotypes (*28*). The reconstructed chromosome 17 variants pinpointed the most likely sequence triggers for the observed rearrangements. The observation that specific locations on the chromosome were repeatedly involved in creating new rearrangements during chromosome degeneration indicates that these locations are fragile sites (*i.e.* sites frequently co-locating with chromosome rearrangements). Three out of four of these fragile sites co-locate with copies of *Styx*. The tendency of *Styx* to trigger chromosomal degeneration is likely widespread within the species, as the same TE is also responsible for an inter-chromosomal rearrangement (*18*). Isolates collected near the geographic center of origin of the pathogen in the Middle East carry no or only few *Styx* copies in the genome and the expansion to higher copy-numbers appears restricted to European genotypes and their descendants (*13*). A separate and likely independent burst of amplification of *Styx* appears to have occurred in the sister species of *Z. tritici*, where no chromosomal rearrangements on chromosome 17 have been described yet (*18*, *29*). The significant consequences for chromosomal integrity could mean that the activity of *Styx* and the presence of fragile sites are under strong selection.

Taken together, our results indicate that specific TE sequences can trigger runaway chromosome degeneration. Non-allelic homologous recombination drives the deleterious rearrangements at the onset of the process with non-disjunction events in subsequent rounds of meiosis. Specific regions on chromosomal segments are preferentially amplified, consistent with patterns observed during degenerative BFB cycles in cancer cell lines (*24*, *30*). BFB was first discovered in maize by McClintock in dicentric chromosomes going through cycles of degeneration (*31*, *32*) and are also known to occur in animals (*33*, *34*) and fungi (*35*, *36*). BFB cycles are initiated via telomere-telomere fusions of chromosomes with degraded or missing telomeres (*37*). The centromeres of the dicentric chromosome are pulled in opposite directions during anaphase, generating a bridge that breaks apart and results in daughter cells with different lengths of the chromosome that lack telomeres (*25*, *33*, *38–40*). BFB cycles tend to be self-perpetuating through repeated fusion and breakage events. At the karyotype level, chromosome 17 undergoes cycles consistent with BFB cycles. However, the likely absence of centromere duplications is inconsistent with classic BFB cycles. We pinpoint an alternative mechanism driving amplification of chromosomal regions in a pattern resembling BFB and involving ectopic recombination. Recombination serves both as the initial trigger to create unstable chromosomes and as the mechanism to maintain the degenerative process. Non-disjunction amplifies the process following the initial meiosis, producing the aberration. We show that large-scale chromosome rearrangements can spontaneously occur in natural pairings of fungal individuals, bearing hallmarks of degenerative processes in somatic cell lines such as those observed in cancers.

## Materials and Methods

### Establishment of a four-generation progeny

To detect how chromosomal rearrangements evolved on chromosome 17, we analyzed four generations of crosses, including two backcrosses of the haploid progeny A66.2 (fig. 1A). The initial cross between parental strains 1E4 and 1A5, and subsequent crosses were described earlier (*7*, *41*). We used previously generated telomere-to-telomere PacBio assembly of parental strains 1E4, 1A5 and 3D7 (*9*, *11*). Parental genome assemblies were validated using high-density genetic maps (*7*, *42*). Crosses were performed by coinfecting wheat leaves with asexual conidia from the parental strains of opposite mating types, according to an established crossing protocol (*43*): Conidia of both parents were sprayed onto wheat plants in equal concentration and incubated outdoors for 40–60 days. Ascospores were isolated over several days by incubating infected wheat leaves on wet filter paper inside Petri dishes. Wheat leaves were covered with upside down Petri dish lids filled with water agar, enabling the capture of vertically ejected ascospores. Ascospores captured on the water agar were left to germinate and inspected for contaminants. Only progeny isolates from single ascospores were selected. Each germinating ascospore was transferred to an individual culture plate for clonal propagation. The mycelium produced by each ascospore was used for DNA extraction. Progeny mycelium was grown in YSB (yeast sucrose broth) liquid medium for 6–7 days at 20°C prior to DNA extraction.

### Chromosome segment PCR assay

In order to assess the presence-absence polymorphism of chromosome 17 segments, we used previously designed PCR assays to amplify ∼500 bp regions of coding sequences at regular intervals along the chromosome 17 of reference strain IPO323 (*7*). Detailed information on primer binding sites in the genome of 1A5 is available in supplementary table S3. PCR reactions were performed in 20 µl volumes with 5–10 ng genomic DNA, 0.5 mM of each primer, 0.25 mM dNTP, 0.6 U Taq polymerase (DreamTaq, Thermo Fisher, Inc.), and the corresponding PCR buffer. In order to avoid false negatives, we included a primer pair for a conserved microsatellite locus in each PCR mix (*44*). The amplification protocol was described in more detail previously (*7*). Successful PCRs produced an additional band that was clearly distinguishable from the PCR product associated with the amplified chromosome region. PCR products were analyzed on agarose gels. Data was visualized using the *R* package *gplots* (https://github.com/talgalili/gplots).

### DNA extraction for PacBio sequencing

Progeny DNA from each cross was extracted using a modified version of the cetyltrimethylammonium bromide (CTAB) DNA extraction protocol developed for plant DNA extractions (*45*). Fungal cultures were grown for 5–7 days in YSB broth and lyophilized overnight. Approximately 60–100 mg of dried material was crushed with a mortar and pestle. The phenol-chloroform-isoamyl alcohol extraction step was performed twice and the washing step three times. In the last step, the DNA pellet was resuspended in 100 µl of sterile water.

### Preparation of fungal material for molecular karyotyping

DNA from intact chromosomes was extracted from conidia from fungal cultures, embedded in agarose gels by the *in situ* digestion of cell walls, using a modified non-protoplasting method (*46*). We included seven *Z. tritici* isolates confirmed with PCR testing to have inherited chromosome 17 from each of the crosses. Isolates were transferred from stocks maintained in glycerol at −80°C to yeast malt agar plates and incubated for 3–4 days in the dark at 18°C. Conidia were then isolated by washing the plates with sterile water and transferring 600–800 μl of suspended conidia to new YMA plates. The plates were again incubated for 2 to 3 days as described above. Conidia were harvested by washing the plates with sterile distilled water and filtered through sterile Miracloth (Calbiochem, La Jolla CA, USA) into 50 ml Falcon tubes. The volume was adjusted to 50 ml by adding more distilled water and the suspension was centrifuged at 3750 rpm at room temperature for 15 min with a clinical centrifuge (Allegra X-12R, Beckman Coulter, Brea CA, USA). The pellets were resuspended in 1–3 ml TE buffer (10 mM Tris-HCL, pH 7.5; 1 mM EDTA, pH 8.0) and vortexed gently. The spore concentration of the solution was calculated using a Thoma haematocytometer cell counter. The 1.5 ml spore suspensions with a concentration between 8×10^7^ to 2×10^8^ spores/ml were transferred to 50 ml Falcon tubes and incubated at 55°C in a water bath for a few minutes. We added 1.5 ml pre-warmed (55°C) low-melting-point agarose prepared in TE Buffer (2% w/v; molecular biology grade, Biofinex, Switzerland). The solution was mixed by gentle pipetting. An aliquot of 500 μl was solidified on ice for approximately 10 min in a pre-cooled plug casting mold (BioRad Laboratories, Switzerland). Agarose plugs were incubated in 15 ml Falcon tubes containing 5 ml of a lysing solution (0.25 M EDTA, pH 8.0, 1.5 mg/mL protease XIV (Sigma, St. Louis MO, USA), 1.0% sodium dodecyl sulfate (Fluka, Switzerland)). Plugs were incubated for ∼24h at 55°C. The lysing solution was changed once after ∼18h and gently mixed every few hours. Plugs with whole chromosomal DNA were washed three times for 15– 20 min in ∼5ml of a 0.1 M EDTA (pH 9.0) solution and then stored in the same solution at 4°C until use.

### Pulsed-field gel electrophoresis

To detect length differences in chromosome 17, pulsed-field gel electrophoresis (PFGE) was performed using a BioRad CHEF II apparatus (BioRad Laboratories, Hercules CA, USA). Chromosomal plugs were placed in the wells of a 1.2% (w/v) agarose gel (Invitrogen, Switzerland) to separate small chromosomes up to 1Mb. Chromosomes were separated at 13°C in 0.56× Tris-borate-EDTA Buffer (*47*) at 200 V with a 60–120 s pulse time gradient for 24–26 h. Gels were stained in ethidium bromide (0.5 mg/ml) for 30 min. De-staining was performed in water for 5–10 min. Photographs were taken under ultraviolet light with a Molecular Imager (Gel Doc XR+, BioRad, Switzerland). As size standards, we used chromosome preparations of *Saccharomyces cerevisiae* (BioRad, Switzerland).

### Southern hybridization for A66.2 and A2.2

Southern blotting and hybridization were performed following standard protocols (*47*). First, hydrolysis was performed in 0.25 M HCl for 30 min, then DNA was transferred onto Amersham Hybond-N+ membranes (GE Healthcare, Switzerland) overnight under alkaline conditions. DNA was heat-fixed onto the membranes at 80°C for 2 h. Membranes were prehybridized overnight with 25 ml of a buffer containing 20% (w/v) SDS, 10% BSA, 0.5 M EDTA (pH 8.0), 1 M sodium phosphate (pH 7.2) and 0.5 ml of sonicated fish sperm solution (Roche Diagnostics, Switzerland). Probes were labeled with ^32^P by nick translation (New England Biolabs, Inc.) following the manufacturer’s instructions. Hybridization was performed overnight at 65°C. Blots were subjected to stringent wash conditions with a first wash in 16 X SSC and 0.1% SDS and a second wash with 0.26 X SSC and 0.1% SDS. Both washes were performed at 60°C. Membranes were exposed to X-ray film (Kodak BioMax MS) for 2 to 3 days at −80°C. We used the same probe as described earlier (supplementary table S4) (*7*).

### PacBio library preparation

PacBio SMRTbell libraries were prepared using 15–31 µg of high-molecular-weight DNA. The libraries were size-selected with an 8 kb cutoff on a BluePippin system (Sage Science, Inc.). After selection, the average fragment length was 15 kb. PacBio sequencing was run on a PacBio RS II instrument or Sequel at the Functional Genomics Center, Zurich, Switzerland using P4/C2 and P6/C4 chemistry, respectively.

### Assembly of chromosome 17 and breakpoint-junction analyses

Each chromosome 17 for all four generations was then assembled (fig. 1A). We mapped the reads to the reference genome 1A5 using minimap2 version 2.17 (*48*) with the parameters --secondary=no -ax map-pb. We compared the coverage of regions of chromosome 17 to the mean coverage of the core chromosomes (1–13) to estimate copy numbers of distinct regions. We then identified breakpoints by analyzing regions with >15 reads that either end or start at a specific position using BEDTools bamtobed version 2.29.2 (*49*) and extracted split reads in this region. We then assembled draft chromosomes by using the structural information from reads showing split alignments, and by joining individual breakpoints (supplementary table S2). Reads were mapped to the assembled chromosomes with minimap2, and we counted the number of reads spanning each established junction point (supplementary table S2). Established chromosome 17 assemblies were error-corrected with Quiver version 2.1.2 (*50*).

### Characterization of sequence features of breakpoints

Transposable elements consensus sequences were annotated on the parent 1A5 chromosome 17 sequence with RepeatMasker version 2.10.0 based on previously described TE family consensus sequences (*13*). The cut-off value was set to 250 and both simple repeats and low-complexity regions were discarded. We used nucmer version 4.0.0rc1 and dotPlotly (https://github.com/tpoorten/dotPlotly/) to visualize the *Styx* TE consensus sequence (*51*). To further classify *Styx*, we analyzed conserved domains in the consensus sequence with online BLASTx on the nonredundant NCBI protein database. We detected four putative coding regions consistent with previous analyses of the TE (*18*). As a proxy for repeat-induced point mutations, we calculated the GC-content for each copy with *geecee* from EMBOSS version 6.6.0 (*52*). To search for microhomology at rearrangement breakpoints, we analyzed tandem repeats within 1000 bp of the breakpoint locations A–E using *mreps* with the parameters -exp 3 and -res 5 to allow for fuzzy detection of degenerate repeats (*53*).

### Genome-wide description of the Styx TE family

We extracted all annotated copies of *Styx* in the 19 reference-quality *Z. tritici* genomes and the sister species (*13*, *20*) with SAMtools faidx version 1.9 (*54*). Multiple sequence alignments of all copies were made with MAFFT version 7.453 using the following parameters: --reorder --localpair --maxiterate 1000 -- nomemsave --leavegappyregion (*55*). We located the putative coding region 4 in the multiple sequence alignment and extracted the sequence with extractalign from EMBOSS. We excluded empty hits (*i.e.* not containing coding regions) with trimAl version 1.4rev15 and sequences with more than 20% of gap sites with SeqKit version 0.11.0 (*56*, *57*). To remove large blocks of gaps and regions that represent rare insertions, we used Gblocks version 0.91b with the parameters: -t=d -b4=5 -b5=h (*58*). We used RAxML version 8.2.12 to create phylogenetic trees in three rounds (*7*): First, we generated 20 ML trees each with a different starting tree and extracted the tree with best likelihood with the following parameters: raxmlHPC-PTHREADS-SSE3 -T 4 -m GTRGAMMA -p 12345 -# 10 --print-identical-sequences. Second, we made a bootstrap search for support values with the following parameters: raxmlHPC-PTHREADS-SSE3 -T 4 -m GTRGAMMA -p 12345 -b 12345 -# 50 --print-identical-sequences. Finally, we drew bipartitions on the best ML tree with bootstrapping: raxmlHPC-PTHREADS-SSE3 -T 4 -m GTRGAMMA -p 12345 -f b -- print-identical-sequences. We imported the tree into R with read.tree from the package treeio version 1.10.0 (https://github.com/YuLab-SMU/treeio), converted it to a tibble object with as.tibble from package tibble version 3.0.1 in tidyverse version 1.3.0 and added the GC-content per sequence as a variable with left_join from the package *dplyr* version 0.8.5 in tidyverse. The tree was then recreated with as.phylo and as.treedata in *treeio* (*59*, *60*). We chose a sequence from *Z. pseudotritici* as an outgroup to root the *Z. tritici* tree. We visualized the tree with GGTree version 2.0.1 (*61*, *62*).

### Repeat-induced point mutation analysis

To detect the impact of RIP in the progeny, we checked for biases of mutations for specific dinucleotides. We mapped PacBio reads to the reference genome 1A5 using minimap2 (*48*) as described above. We performed SNP calling with the software Longshot version 0.4.0 and included a minimum mapping quality cutoff of -q 30 (*63*). All SNPs with a “dn” tag, indicating mapping issues, were removed. We allowed for 20% errors in the uncorrected PacBio reads. We expected the alternate allele at polymorphic sites to be shared by ≥80% of the mapped reads for single-copy regions, shared by ≥40% of the reads in regions with evidence for duplications (based on read coverage), shared by ≥27%, ≥20% and ≥16% of the reads in regions with three, four and five copies (based on read coverage), respectively. We also excluded regions with ≤50% or ≥150% of the expected read coverage for further analyses to reduce erroneous variant calls due to inconsistent read mapping.

## Data availability

The genome assembly and annotation for 1E4, 1A5, and 3D7 genome are available at the European Nucleotide Archive (http://www.ebi.ac.uk/ena) under accession numbers PRJEB15648, PRJEB20900 and PRJEB14341. Progeny genomes are available under the project PRJNA645795.

## Author contributions

SF and DC conceived the study; SF and UO performed analyses; BAM and DC supervised the work and provided funding; SF, UO and DC wrote the manuscript with input from all co-authors.

## Acknowledgements

We are grateful for technical support in the laboratory by Marcello Zala. Thomas Badet provided advice on genome sequence analyses.

## Supplementary Materials

Supplementary Figures S1–S12 and Supplementary Tables S1–S7.

