## Supplementary Figures for "Recurrent chromosome destabilization through repeat-mediated rearrangements in a fungal pathogen"

**This PDF file includes:**

Figs. S1 to S12

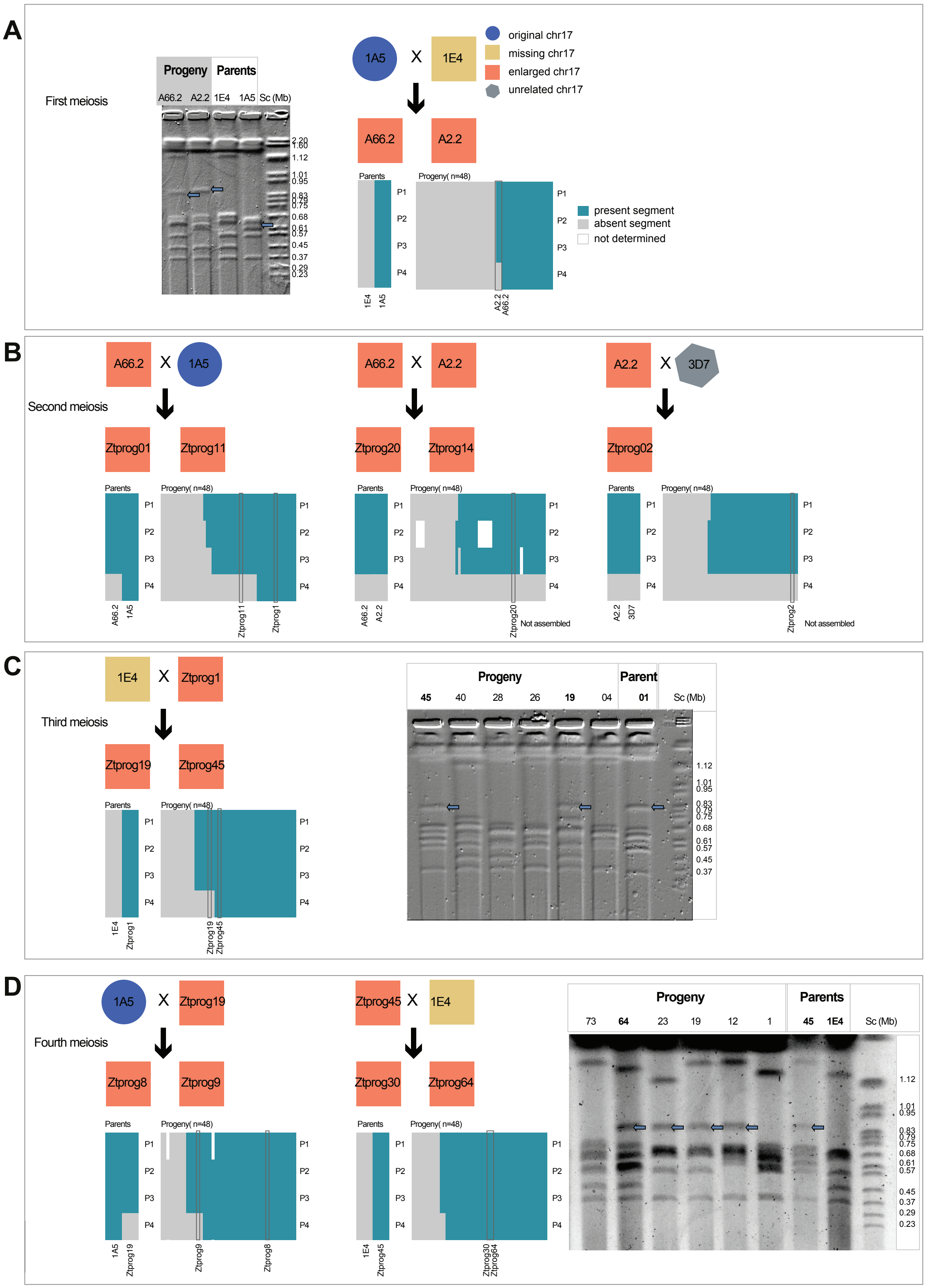
A
First meiosis
Progeny
Parents
A66.2 A2.2 1E4 1A5 Sc (Mb)
1A5 X 1E4
original chr17
missing chr17
enlarged chr17
unrelated chr17
A66.2 A2.2
present segment
absent segment
not determined
B
Second meiosis
A66.2 X 1A5
Ztprog01 Ztprog11
A66.2 X A2.2
Ztprog20 Ztprog14
A2.2 X 3D7
Ztprog02
Not assembled
C
Third meiosis
1E4 X Ztprog1
Ztprog19 Ztprog45
Progeny
Parent
45 40 28 26 19 04 01 Sc (Mb)
D
Fourth meiosis
1A5 X Ztprog19
Ztprog8 Ztprog9
Ztprog45 X 1E4
Ztprog30 Ztprog64
Progeny
Parents
73 64 23 19 12 1 45 1E4 Sc (Mb)

**Supplementary Figure 1: Chromosome pedigree of parents and progeny of four rounds of meiosis (A-D).** The colors of the block indicate whether the isolate carries either an original, enlarged or no chromosome 17 according to pulsed-field gel electrophoresis (PFGE) analysis. Barplots below each cross summarize the presence (turquoise) or absence (grey) of four ~500 bp segments of coding sequences at regular intervals along the chromosome 17 (P1-P4) assayed by PCR  in parents and progeny (n=48) from each cross. Electrophoretic karyotype diversity of chromosome 17 among a selection of progeny from each cross are shown along parental karyotypes. The size marker is *Saccharomyces cerevisiae* chromosomes (Sc). Arrows indicate the most likely band representing chromosome 17.

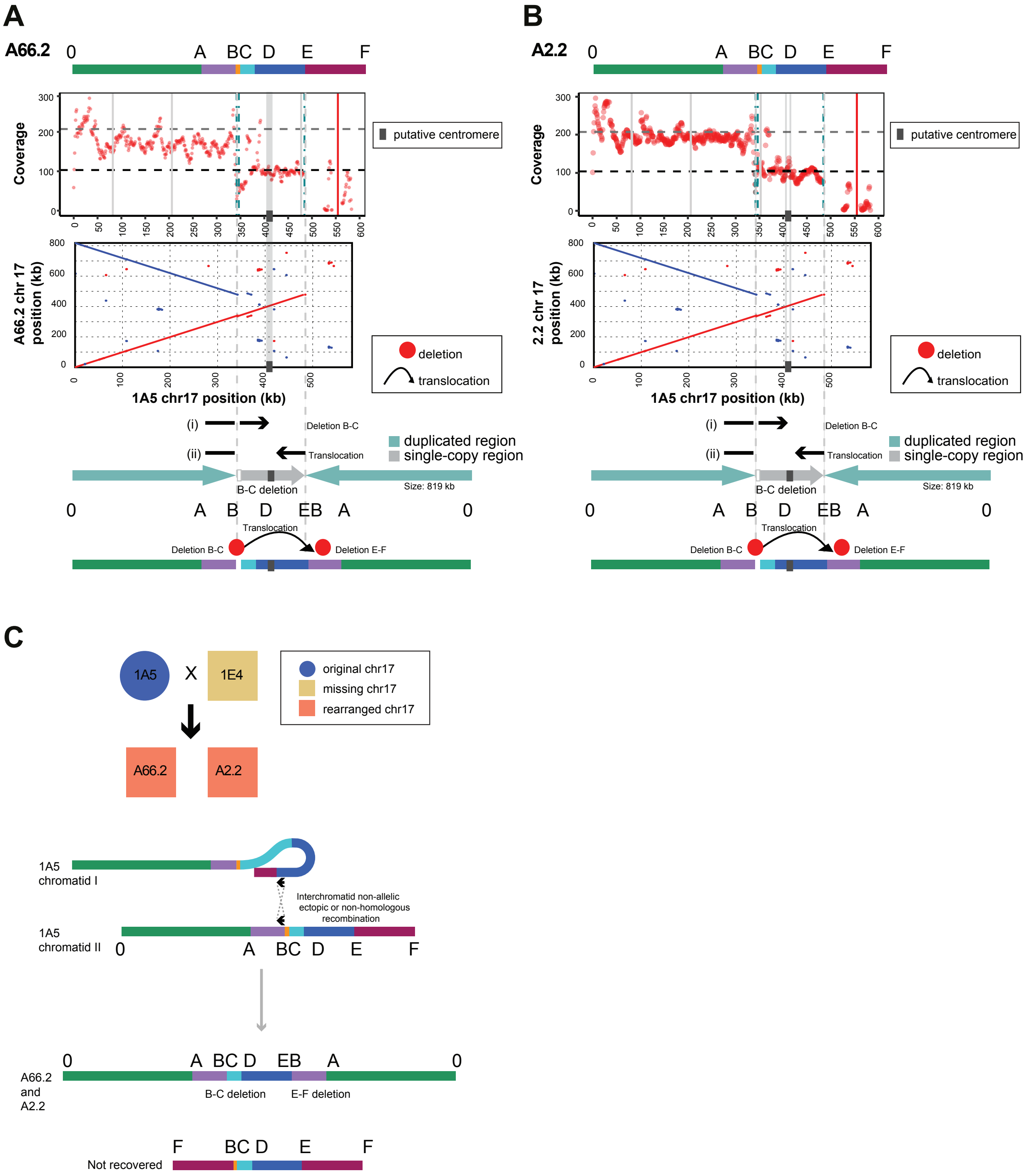

**Supplementary Figure 2: Generation of the enlarged chromosome 17 in A66.2 and A2.2.** (A) The coverage and breakpoints of the progeny A66.2 and A2.2 reads mapped to the parent 1A5, horizontal dashed lines indicate the mean coverage of the core chromosomes (black) and in grey two times the mean core chromosome coverage. Red dots indicate the mean coverage in 1 kb windows (regions with excessive, >300 x coverage are removed). Vertical dashed lines indicate the chromosomal breakpoints identified from mapped reads (B and E, turquoise and continuing in grey). Solid vertical lines indicate the positions of loci amplified by PCR (grey=positive and red=negative). Dotplots of the assembled chromosome for progeny A66.2 and A2.2 compared to the parent 1A5. Inverted regions are indicated in blue. (i) and (ii) show the location and continuation of split reads and below is a schematic of the resulting enlarged chromosome in the progeny. (B) Pulsed-field gel electrophoresis of the parental chromosomes (1E4 and 1A5) and the progeny A2.2 and A66.2, showing the enlarged chromosome 17 (arrows; Croll et al., 2013). (C) Schematic representation of the breakpoints and rearrangements between the two chromatids of 1A5 that generated the enlarged chromosome 17 recovered in the progeny A66.2 and A2.2. The other product that was not recovered is also indicated.

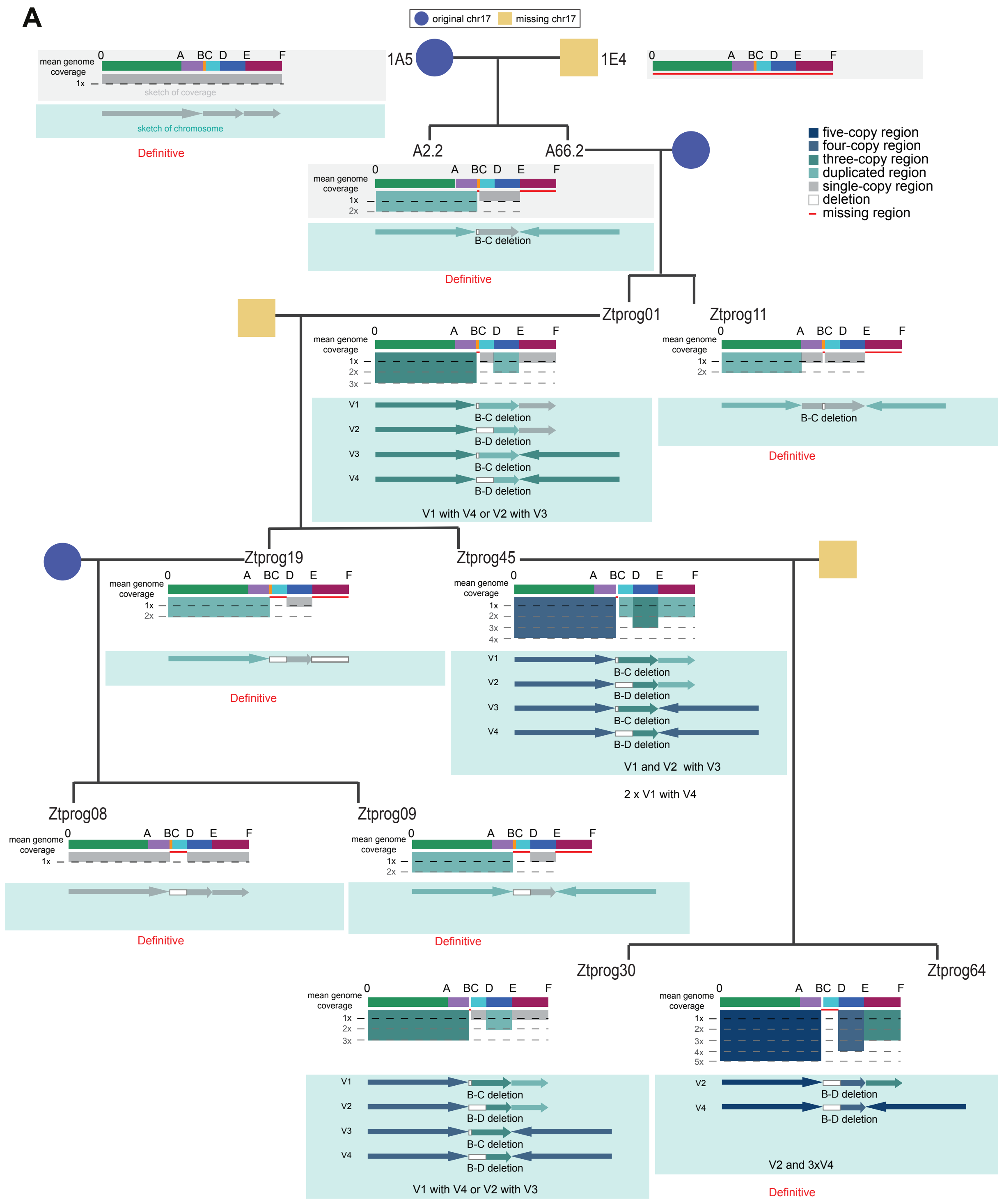


**Supplementary figure 3: Reconstruction of chromosome 17 variants based on coverage analysis through four generations of the pedigree.** Schematic of the coverage of reads mapped to the parent 1A5 chromosome 17 in each progeny. Coverage is shown relative to the mean genome coverage of the core chromosomes. The expected variants present in the progeny are also indicated.

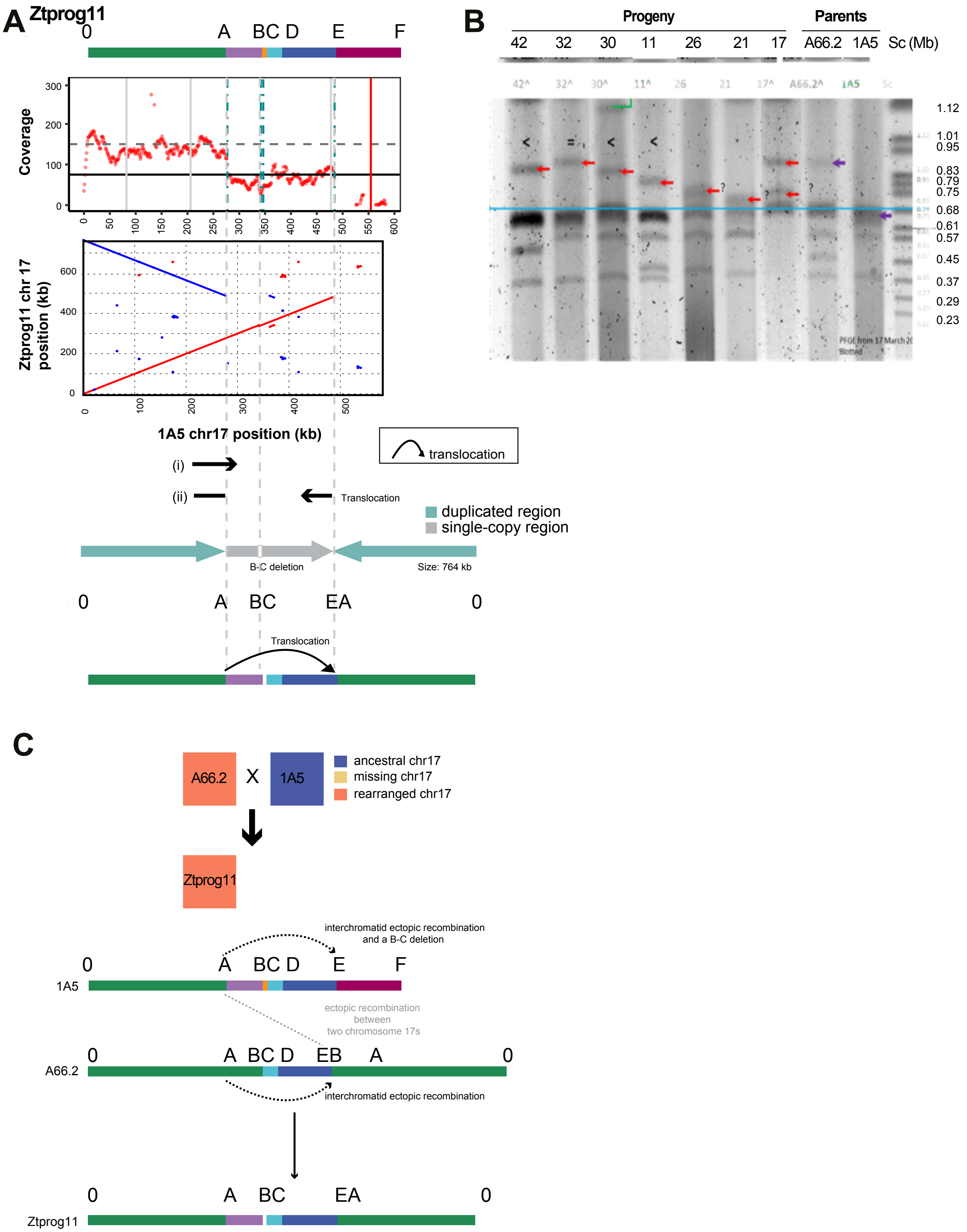


**Supplementary Figure 4: Generation of the rearranged chromosome 17 in Ztprog11.** (A) The coverage and breakpoints of the progeny Ztprog11 reads mapped to the parent 1A5, horizontal dashed lines indicate the mean coverage of the core chromosomes (black) and in grey two times the mean core chromosome coverage. Red dots indicate the mean coverage in 1 kb windows (regions with excessive, >300 x coverage are removed). Vertical dashed lines indicate the chromosomal breakpoints identified from mapped reads (A,B and E, turquoise and continuing in grey). Solid vertical lines indicate the positions of loci amplified by PCR (grey=positive and red=negative). Dotplots of the assembled chromosome 17 for progeny Ztprog11 compared to the parent 1A5. Inverted regions are indicated in blue. (i) and (ii) show the location and continuation of split reads and below is a schematic of the resulting enlarged chromosome in the progeny. (B) Pulsed-field gel electrophoresis of the parental chromosomes (A66.2 and 1A5) and the progeny Ztprog11, showing the enlarged chromosome. (C) Schematic representation of the breakpoints and rearrangements between the chromosomes in A66.2 and 1A5 (inter or intrachromosomal rearrangements) that may have resulted in the chromosome in the progeny Ztprog11.

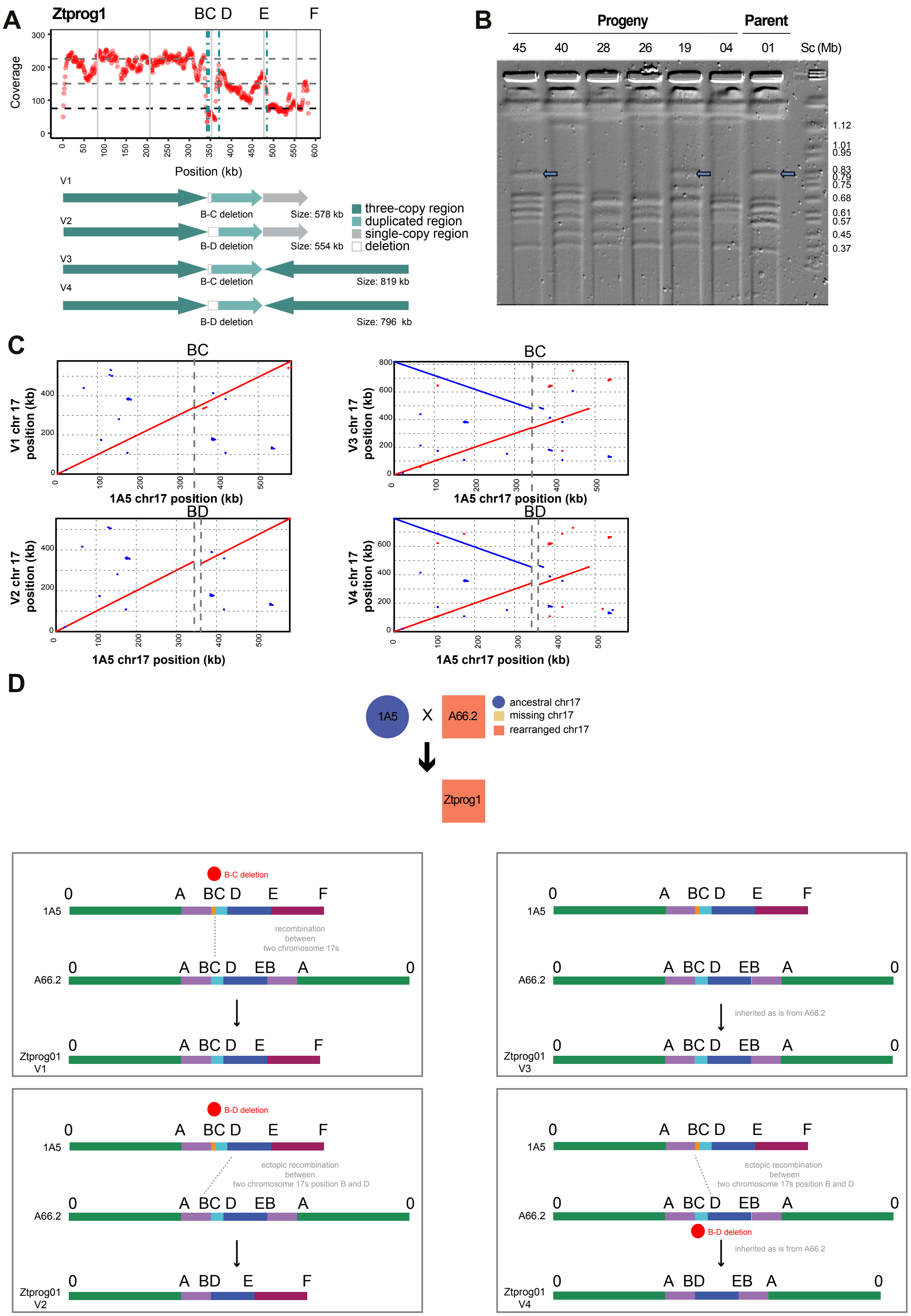
A
Ztprog1
BC D
E
F
Coverage
Position (kb)
V1
B-C deletion
Size: 578 kb
V2
B-D deletion
Size: 554 kb
V3
B-C deletion
Size: 819 kb
V4
B-D deletion
Size: 796 kb
three-copy region
duplicated region
single-copy region
deletion
B
Progeny
Parent
45 40 28 26 19 04 01 Sc (Mb)
1.12
1.01
0.95
0.83
0.79
0.75
0.68
0.61
0.57
0.45
0.37
C
BC
BD
V1 chr 17 position (kb)
V2 chr 17 position (kb)
V3 chr 17 position (kb)
V4 chr 17 position (kb)
1A5 chr17 position (kb)
D
1A5
X
A66.2
ancestral chr17
missing chr17
rearranged chr17
Ztprog1
B-C deletion
B-D deletion
recombination between two chromosome 17s
ectopic recombination between two chromosome 17s position B and D
inherited as is from A66.2
0 A BC D E F
0 A BC D EB A 0
Ztprog01 V1
Ztprog01 V2
Ztprog01 V3
Ztprog01 V4
A66.2
0 A BD E F
0 A BD EB A 0

**Supplementary Figure 5: Generation of the rearranged chromosome 17 in Ztprog01.** (A) The coverage and breakpoints of the progeny Ztprog01 reads mapped to the parent 1A5, horizontal dashed lines indicate the mean coverage of the core chromosomes (black) and in grey two and three times the mean core chromosome coverage. Red dots indicate the mean coverage in 1 kb windows (regions with excessive, >300 x coverage are removed). Vertical dashed lines indicate the chromosomal breakpoints identified from mapped reads (turquoise). Solid vertical lines indicate the positions of loci amplified by PCR (grey=positive and red=negative). Below each coverage plot: variants of chromosome 17 that are likely to be in this progeny isolate. (B) Pulsed-field gel electrophoresis of the chromosomes of progeny Ztprog01, showing the enlarged chromosome. (C) Dotplots of the assembled chromosome variants (V1-V4) for progeny Ztprog01 compared to the parent 1A5. Inverted regions are indicated in blue. Vertical dashed lines indicate the chromosomal breakpoints identified from mapped reads (grey). (D) Schematic representation of the breakpoints and rearrangements between the chromosomes in A66.2 and 1A5 (inter- or intrachromosomal rearrangements) that may have resulted in the chromosome variant in the progeny Ztprog01.

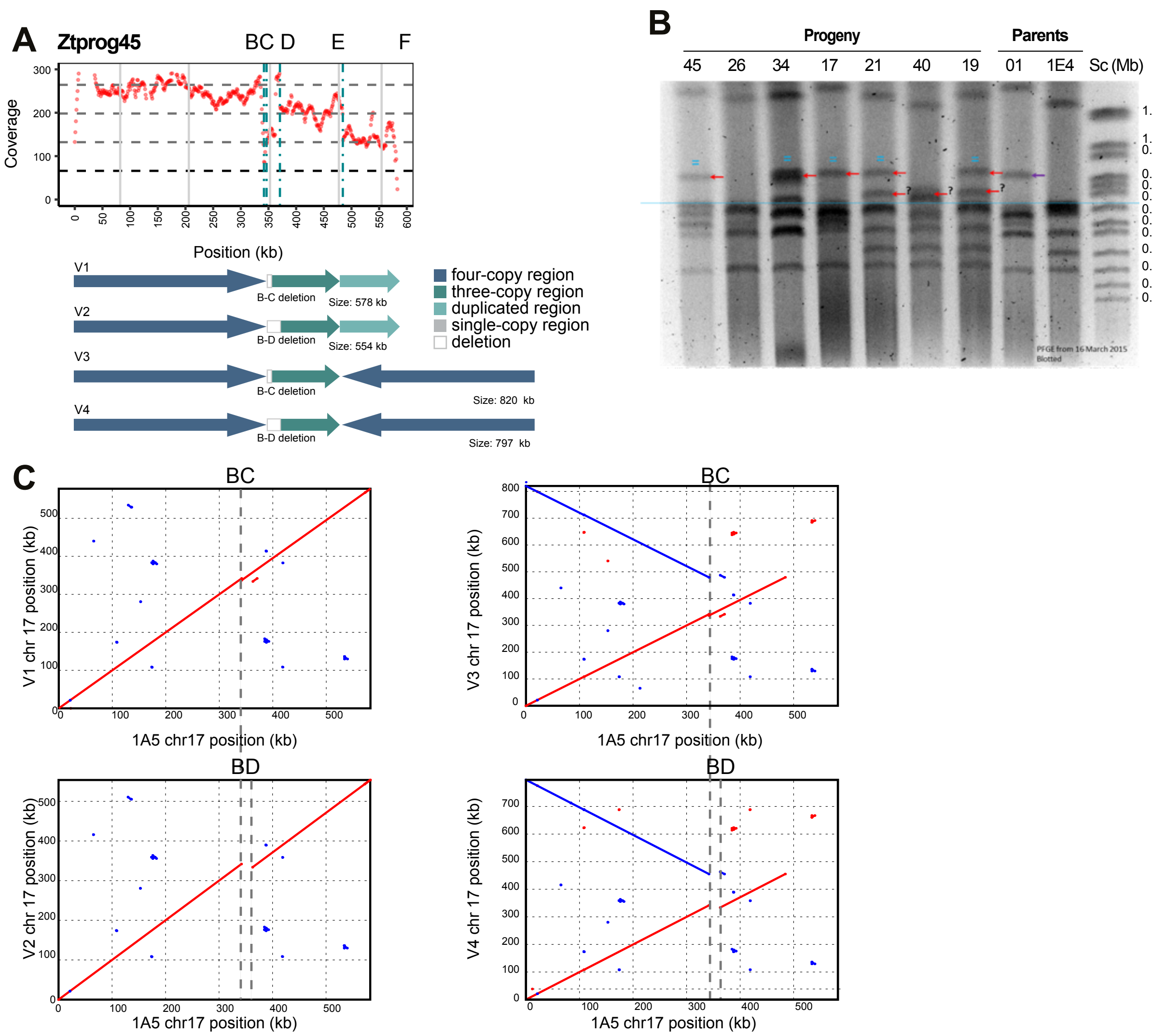


**Supplementary Figure 6: Generation of the rearranged chromosome 17 in Ztprog45.** (A) The coverage and breakpoints of the progeny Ztprog45 reads mapped to the parent 1A5, horizontal dashed lines indicate the mean coverage of the core chromosomes (black) and in grey two, three and four times the mean core chromosome coverage. Red dots indicate the mean coverage in 1 kb windows (regions with excessive, >300 x coverage are removed). Vertical dashed lines indicate the chromosomal breakpoints identified from mapped reads (turquoise). Solid vertical lines indicate the positions of loci amplified by PCR (grey=positive and red=negative). Below each coverage plot: variants of chromosome 17 that are likely to be in this progeny isolate. (B) Pulsed-field gel electrophoresis of the parental chromosomes of 1E4 and Ztprog01 and the progeny progeny Ztprog45, showing the enlarged chromosome. (C) Dotplots of the assembled chromosome variants (V1-V4) for progeny Ztprog45 compared to the parent 1A5. Inverted regions are indicated in blue. Vertical dashed lines indicate the chromosomal breakpoints identified from mapped reads (grey).

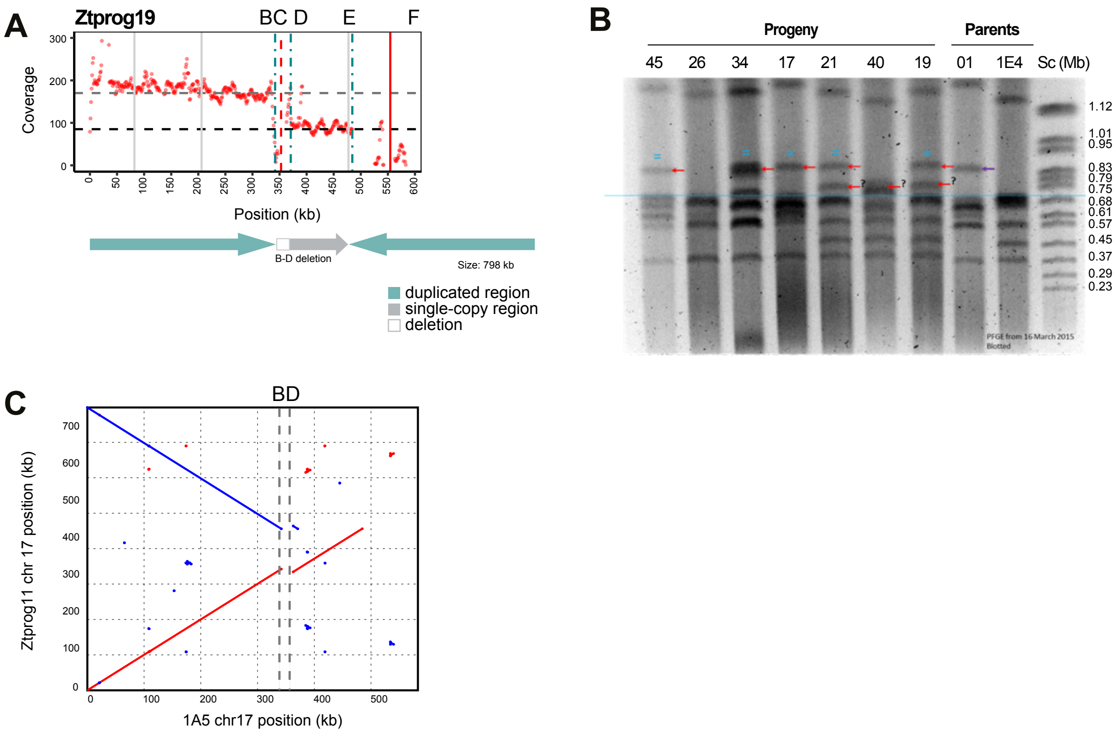


**Supplementary Figure 7: Generation of the rearranged chromosome 17 in Ztprog19.** (A) The coverage and breakpoints of the progeny Ztprog19 reads mapped to the parent 1A5, horizontal dashed lines indicate the mean coverage of the core chromosomes (black) and in grey two times the mean core chromosome coverage. Red dots indicate the mean coverage in 1 kb windows (regions with excessive, >300 x coverage are removed). Vertical dashed lines indicate the chromosomal breakpoints identified from mapped reads (turquoise). Solid vertical lines indicate the positions of loci amplified by PCR (grey=positive and red=negative). Below each coverage plot: variant of chromosome 17 that is likely to be in this progeny isolate. (B) Pulsed-field gel electrophoresis of the parental chromosomes of 1E4 and Ztprog01 and the progeny progeny Ztprog19, showing the enlarged chromosome. (C) Dotplots of the assembled chromosome for progeny Ztprog19 compared to the parent 1A5. Inverted regions are indicated in blue. Vertical dashed lines indicate the chromosomal breakpoints identified from mapped reads (grey).

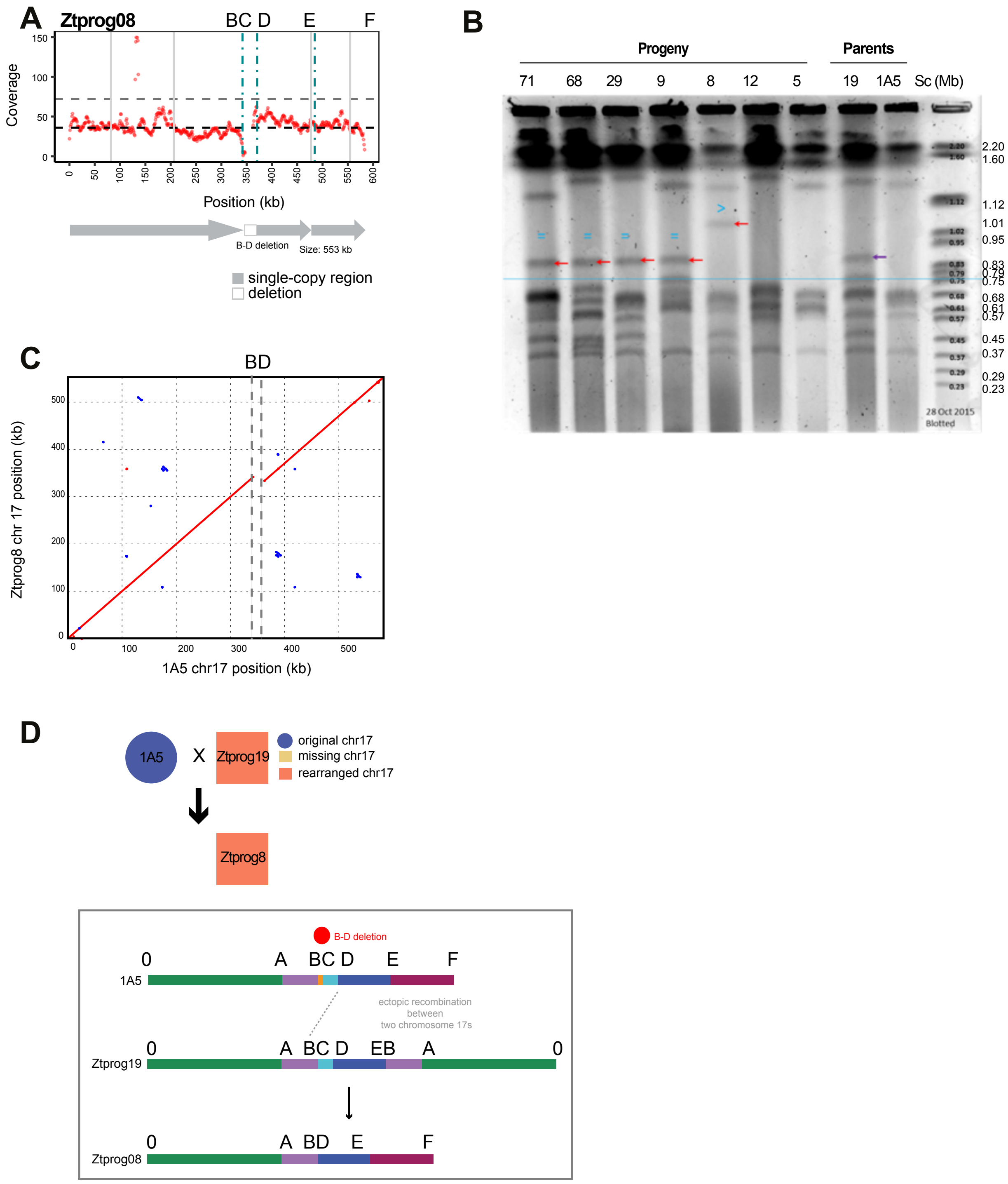


**Supplementary Figure 8: Generation of the rearranged chromosome 17 in Ztprog08.** (A) The coverage and breakpoints of the progeny Ztprog08 reads mapped to the parent 1A5, horizontal dashed lines indicate the mean coverage of the core chromosomes (black) and in grey two times the mean core chromosome coverage. Red dots indicate the mean coverage in 1 kb windows (regions with excessive, >150 x coverage are removed). Vertical dashed lines indicate the chromosomal breakpoints identified from mapped reads (turquoise). Solid vertical lines indicate the positions of loci amplified by PCR (grey=positive and red=negative). Below each coverage plot: variant of chromosome 17 that is likely to be in this progeny isolate. (B) Pulsed-field gel electrophoresis of the parental chromosomes of 1A5 and Ztprog19 and the progeny progeny Ztprog08, showing the enlarged chromosome. (C)

Dotplots of the assembled chromosome for progeny Ztprog08 compared to the parent 1A5. Vertical dashed lines indicate the chromosomal breakpoints identified from mapped reads (grey).

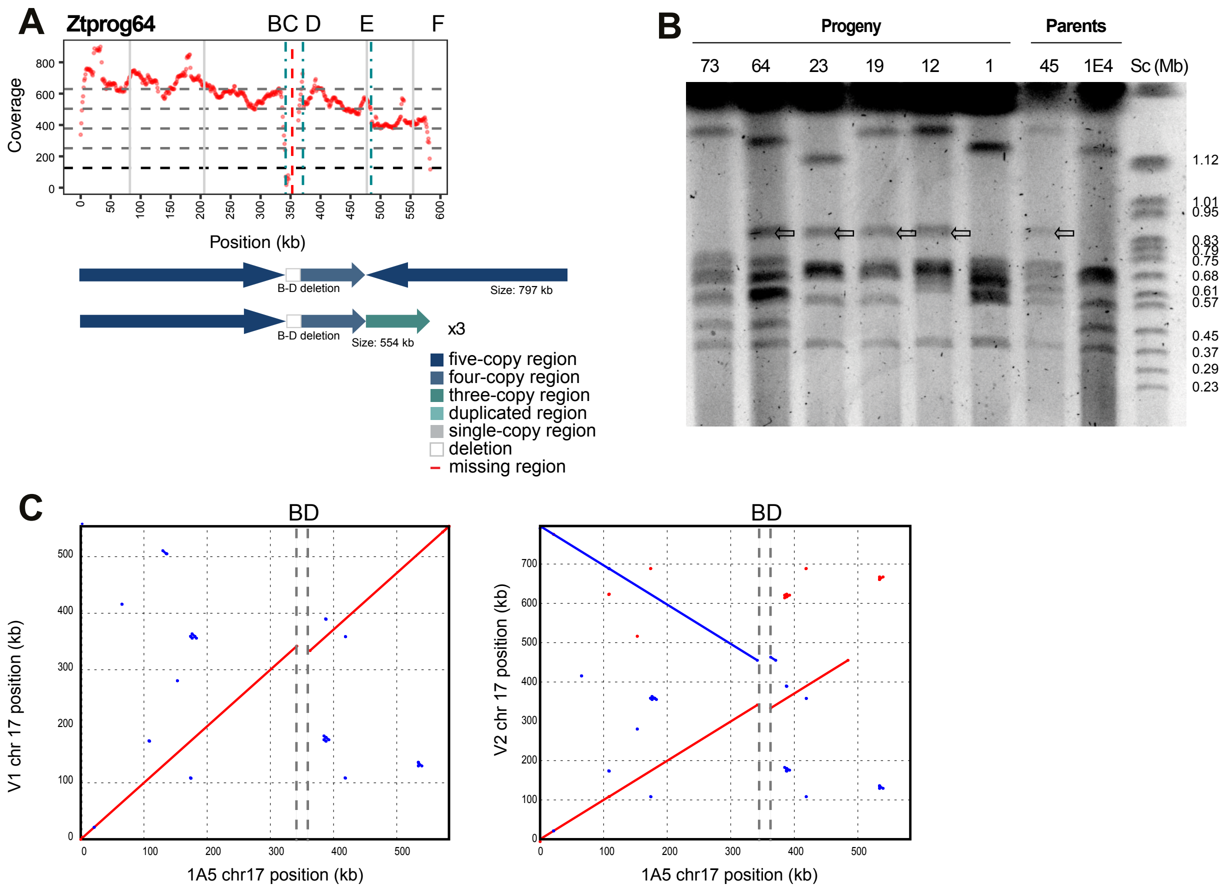


**Supplementary Figure 9: Generation of the rearranged chromosome 17 in Ztprog64.** (A) The coverage and breakpoints of the progeny Ztprog64 reads mapped to the parent 1A5, horizontal dashed lines indicate the mean coverage of the core chromosomes (black) and in grey two, three, four and five times the mean core chromosome coverage. Red dots indicate the mean coverage in 1 kb windows (regions with excessive, >900 x coverage are removed). Vertical dashed lines indicate the chromosomal breakpoints identified from mapped reads (turquoise). Solid vertical lines indicate the positions of loci amplified by PCR (grey=positive and red=negative). Below each coverage plot: variants of chromosome 17 that are likely to be in this progeny isolate. (B) Pulsed-field gel electrophoresis of the parental chromosomes of 1E4 and Ztprog45 and the progeny progeny Ztprog64, showing the enlarged chromosome. (C) Dotplots of the assembled chromosome variants (V1-V2) for progeny Ztprog64 compared to the parent 1A5. Inverted regions are indicated in blue. Vertical dashed lines indicate the chromosomal breakpoints identified from mapped reads (grey).

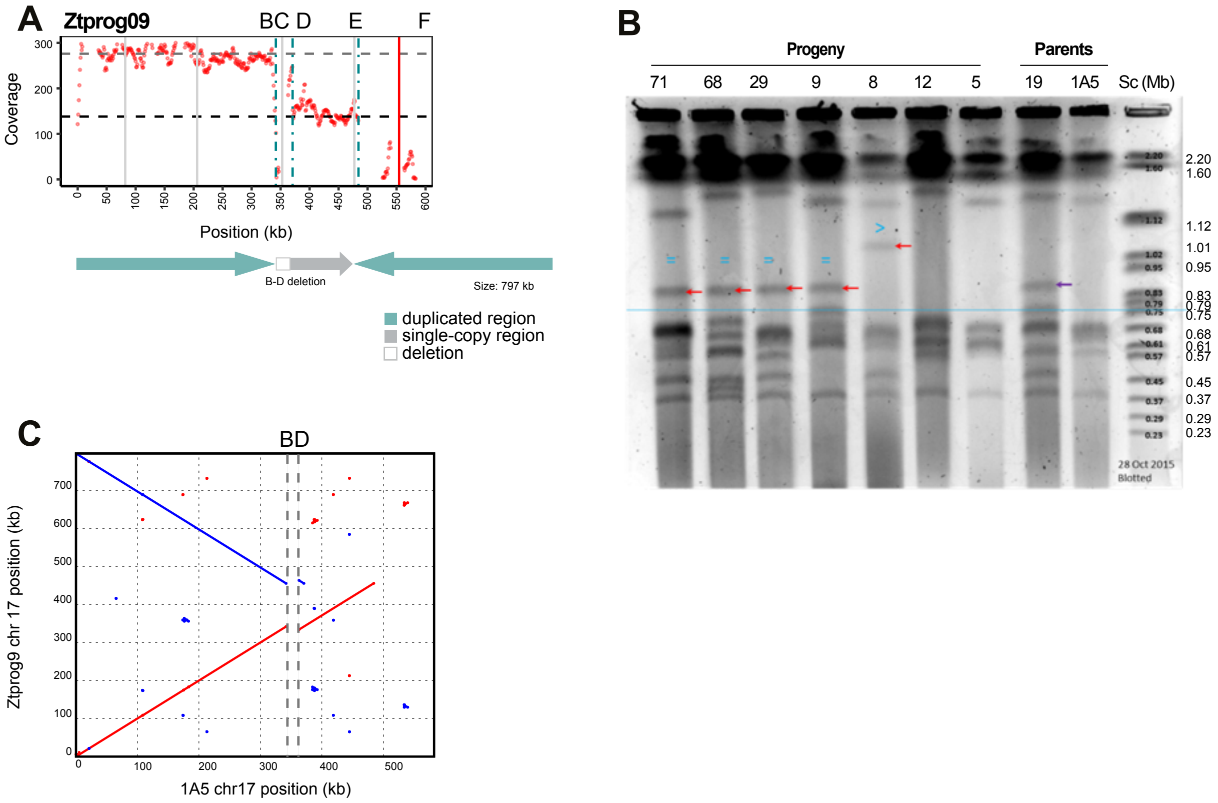


**Supplementary Figure 10: Generation of the rearranged chromosome 17 in Ztprog09.** (A) The coverage and breakpoints of the progeny Ztprog09 reads mapped to the parent 1A5, horizontal dashed lines indicate the mean coverage of the core chromosomes (black) and in grey two times the mean core chromosome coverage. Red dots indicate the mean coverage in 1 kb windows (regions with excessive, >300 x coverage are removed). Vertical dashed lines indicate the chromosomal breakpoints identified from mapped reads (turquoise). Solid vertical lines indicate the positions of loci amplified by PCR (grey=positive and red=negative). Below the coverage plot: variant of chromosome 17 that is likely to be in this progeny isolate. (B) Pulsed-field gel electrophoresis of the parental chromosomes of 1A5 and Ztprog19 and the progeny progeny Ztprog09, showing the enlarged chromosome. (C) Dotplots of the assembled chromosome for progeny Ztprog09 compared to the parent 1A5. Vertical dashed lines indicate the chromosomal breakpoints identified from mapped reads (grey).

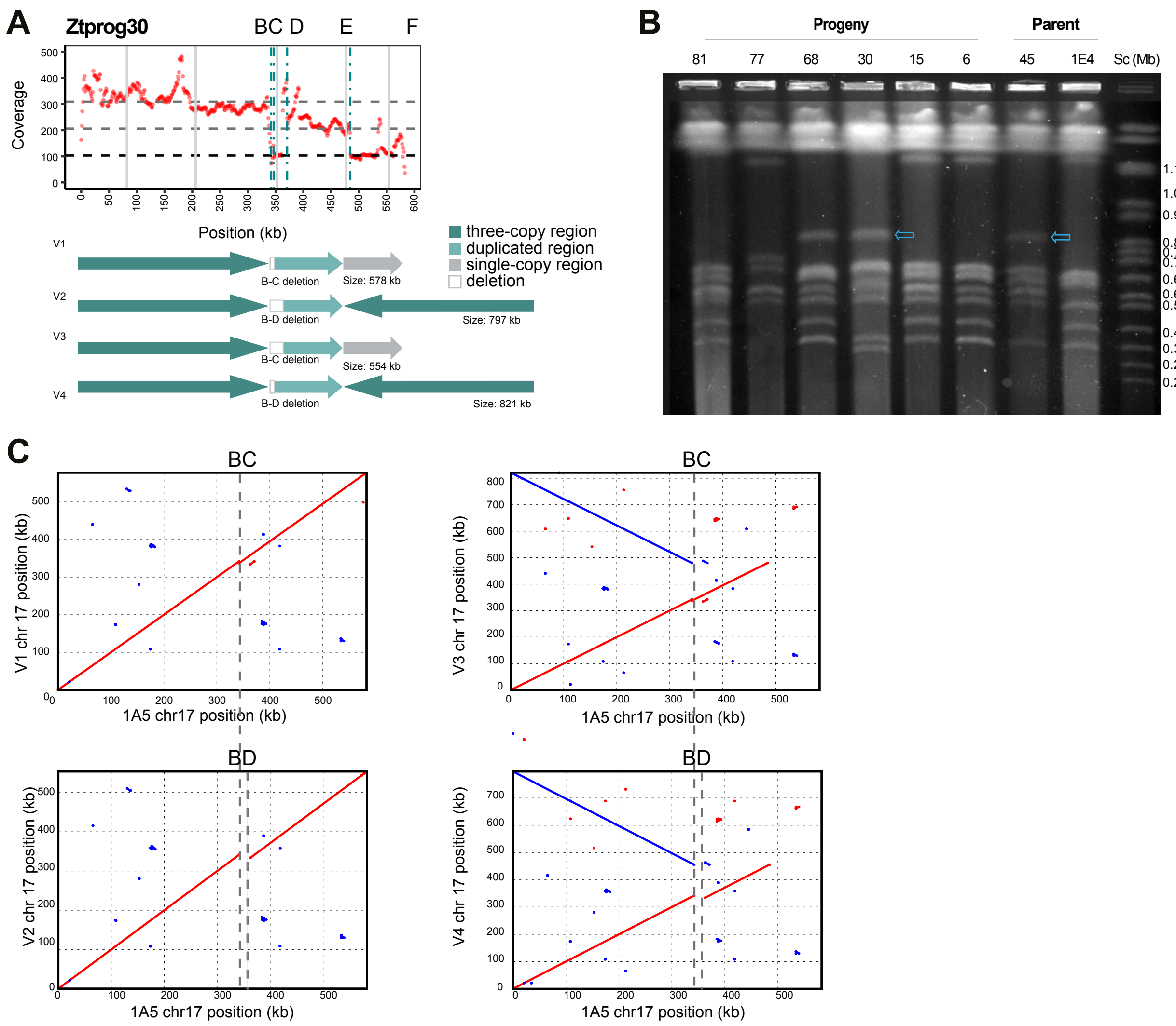


**Supplementary Figure 11: Generation of the rearranged chromosome 17 in Ztprog30.** (A) The coverage and breakpoints of the progeny Ztprog30 reads mapped to the parent 1A5, horizontal dashed lines indicate the mean coverage of the core chromosomes (black) and in grey two and three times the mean core chromosome coverage. Red dots indicate the mean coverage in 1 kb windows (regions with excessive, >500 x coverage are removed). Vertical dashed lines indicate the chromosomal breakpoints identified from mapped reads (turquoise). Solid vertical lines indicate the positions of loci amplified by PCR (grey=positive and red=negative). Below each coverage plot: variants of chromosome 17 that are likely to be in this progeny isolate. (B) Pulsed-field gel electrophoresis of the parental chromosomes of 1E4 and Ztprog45 and the progeny progeny Ztprog30, showing the rearranged chromosome. (C) Dotplots of the assembled chromosome variants (V1-V4) for progeny Ztprog30 compared to the parent 1A5. Inverted regions are indicated in blue. Vertical dashed lines indicate the chromosomal breakpoints identified from mapped reads (grey).

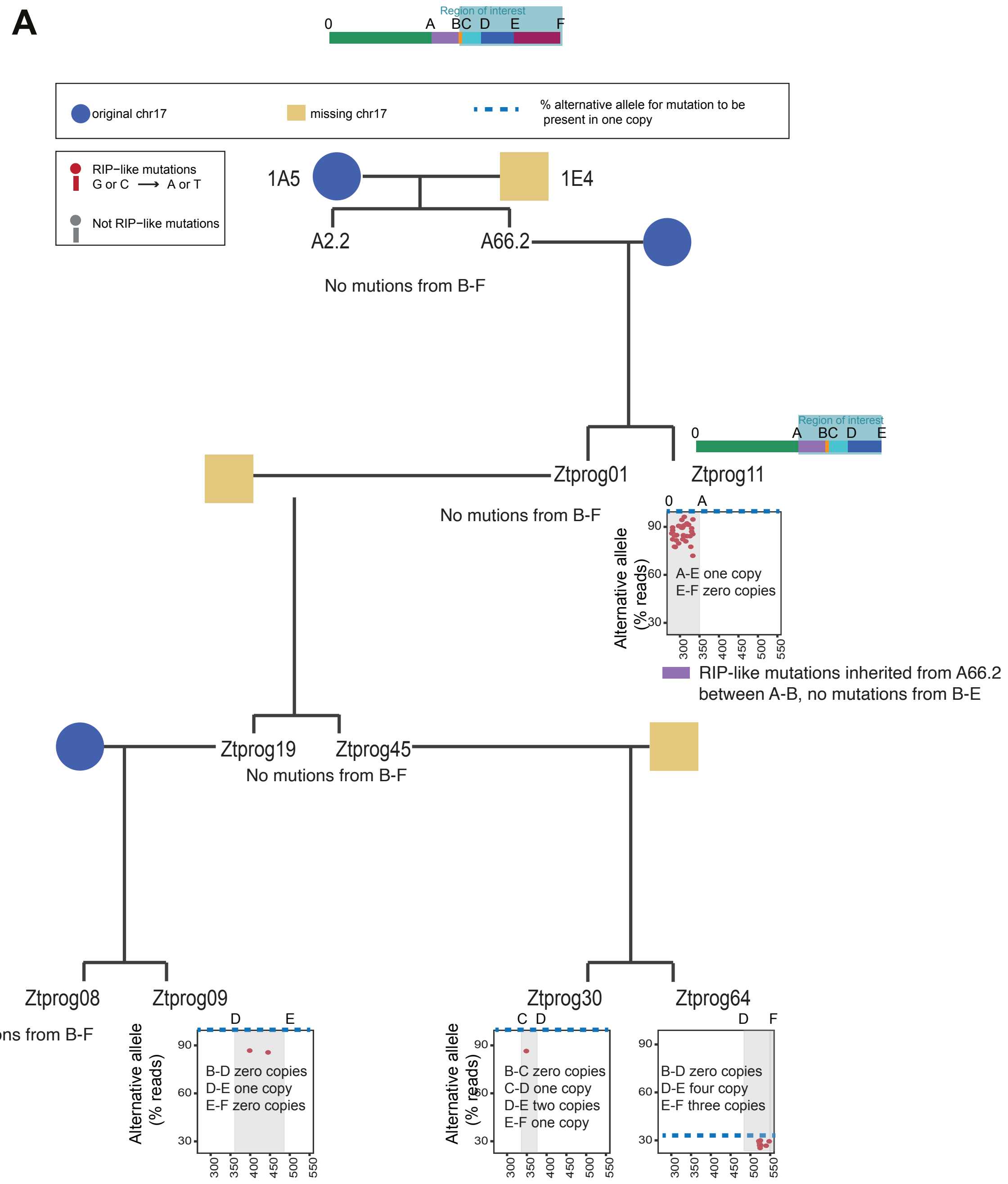


**Supplementary Figure 12: Characterization of repeat-induced point mutation (RIP) working on other sequences.** Dotplots showing the percentage alternative allele in reads mapped to the reference in regions not included in fig. 4 (B-F and A-F for Ztprog11). Copy numbers of the region is indicated and the expected % reads for the mutation to be present in at least one copy of the region (blue dashed line).
